# Single-molecule 3D orientation imaging reveals nanoscale compositional heterogeneity in lipid membranes

**DOI:** 10.1101/2020.05.12.091421

**Authors:** Jin Lu, Hesam Mazidi, Tianben Ding, Oumeng Zhang, Matthew D. Lew

## Abstract

In soft matter, thermal energy causes molecules to continuously translate and rotate, even in crowded environments, impacting the spatial organization and function of most molecular assemblies, such as lipid membranes. Directly measuring the orientation and spatial organization of large collections (>3000 molecules/μm^2^) of single molecules with nanoscale resolution remains elusive. We present SMOLM, single-molecule orientation localization microscopy, to directly measure the orientation spectra (3D orientation plus “wobble”) of lipophilic probes transiently bound to lipid membranes, revealing that Nile red’s (NR) orientation spectra are extremely sensitive to membrane chemical composition. SMOLM images resolve nanodomains and enzyme-induced compositional heterogeneity within membranes, where NR within liquid-ordered vs. liquid-disordered domains shows a ~4° difference in polar angle and a ~0.3π sr difference in wobble angle. As a new type of imaging spectroscopy, SMOLM exposes the organizational and functional dynamics of lipid-lipid, lipid-protein, and lipid-dye interactions with single-molecule, nanoscale resolution.

## Introduction

Tracking a molecule’s 3D position and orientation (and associated translational and rotational motions) within soft matter is critical for understanding the intrinsically heterogeneous and complex interactions of its various components across length scales. In living cells, the organization of many biomolecular assemblies, such as lipid membranes, chromosomes, and cytoskeletal proteins,^1–4^ ensure the proper functioning of all cellular compartments. Molecular organization also significantly impacts the nanoscale morphology of supramolecular structures,^5^ the physical and mechanical properties of polymers,^6^ and carrier mobility in light-emitting diodes.^7^

Molecular orientations are commonly inferred from an order parameter determined via X-ray diffraction,^8^ infrared spectroscopy,^9^ nuclear magnetic resonance (NMR),^10^ Raman spectroscopy,^11^ sum frequency generation spectroscopy,^12^ and fluorescence microscopy.^13^ However, the order parameter is an ensemble average taken and cannot unambiguously determine the 3D orientation of a single molecule (SM).^14^ Spectrally-resolved SM localization microscopy (SMLM)^15–17^ maps the local polarity or hydrophobicity of protein aggregates and subcellular structures,^17,18^ and fluorescence lifetime imaging identifies sub-resolution lipid domains in the plasma membrane.^19^ However, these approaches require specific environment-sensitive fluorescent probes (e.g., Nile red,^20^ Laurdan,^21^ and 3-hydroxyflavone derivatives^22^) whose fluorescence spectra (intensities) or lifetimes are sensitive to local environment.

Alternatively, the orientation and motion of any fluorescent probe are directly influenced by surrounding molecules. Therefore, imaging the 3D orientation and wobble of SMs, which we term as “orientation spectra” in this work, offers an alternative strategy for sensing molecular interactions using any SMLM-compatible fluorescent dye. Numerous imaging technologies can characterize SM orientation with varying degrees of sensitivity and resolution,^23–26^ but to our knowledge, no demonstrated technique is capable of imaging the positions and 3D orientations of large collections of molecules (>3000 molecules/μm^2^) with single-molecule sensitivity and sufficient spatiotemporal resolution to visualize, for example, dynamic remodeling of a lipid bilayer. To address these limitations, we have SMOLM, single-molecule orientation localization microscopy, comprising 1) an engineered point spread function (PSF) that efficiently encodes the 3D orientation and wobble of dipole-like emitters^27^ into fluorescence images and 2) a maximum likelihood estimator with joint-sparse regularization for estimating molecular position, orientation, and wobble from those images.^28,29^ This combination of hardware and software is critical for resolving molecular positions and orientations robustly; otherwise, neighboring molecules, wobbling molecules, and translationally diffusing molecules could be confused with one another.

In this paper, we report the practical application of SMOLM to measure the orientation spectra of single lipophilic dyes in lipid membranes and observe that their characteristic orientation spectra are determined by their chemical structures and surrounding lipid environment. In particular, we discover that the orientation spectra of Nile red (NR), a classic solvatochromic dye,^30^ are extremely sensitive to the composition and packing of lipid membranes. To achieve high sampling density for SMOLM, we apply the PAINT (points accumulation for imaging in nanoscale topography) blinking mechanism,^31^ in which certain lipophilic dyes exhibit fluorescence solely while in a non-polar environment. We thereby resolve nanoscale lipid domains with resolution beyond the diffraction limit and monitor *in-situ* lipid compositional changes induced by low doses of sphingomyelinase. SMOLM imaging clearly shows its potential to resolve interactions between various lipid molecules, enzymes, and fluorescent probes with detail that has never been achieved previously.

## Results

### Resolving the orientation spectra of single molecules

We first utilize SMOLM to image lipophilic dyes with known orientation spectra within supported lipid bilayers (SLBs). Orientation spectra, which are characteristics of the molecules, may be inferred from angular emission spectra and polarization spectra, which are characteristics of the detected photons. Most organic fluorescent probes are well-approximated as oscillating electric dipoles.^13^ We model the orientation of each emission dipole using an average polar angle (θ) and azimuthal angle (ϕ), plus a uniform wobble (rotational diffusion) within a hard-edged cone (solid angle, Ω) in 3D (Fig. 1a); the parameters can be readily adapted for other types of rotational diffusion.^32,33^ An SM’s orientation and wobble is affected by how its molecular structure interacts with its local environment. DiI, for example, bears two long hydrocarbon chains that incorporate into the nonpolar core of a lipid bilayer, while its chromophore headgroup resides in the charged polar region of the bilayer (Fig. 1b). We captured images of single DiI molecules in x- and y-polarized emission channels using the orientation-sensitive Tri-spot PSF^27^ (Fig. 1c(i)(ii)) and estimated molecular orientation and wobble using a maximum-likelihood algorithm developed by our group.^28,29^ Our results (Fig. 1c(iii)) indicate that in 1,2-dipalmitoyl-sn-glycero-3-phosphocholine (DPPC) SLBs, most DiI molecules exhibit large polar angles (θ=73.6±21.2°, median±std) corresponding to an orientation approximately parallel to the plane of the coverslip,^34^ which is corroborated by molecular dynamics simulations.^35^ The solid “wobble” angles of DiI are small (Ω=0.21π±0.41π sr), implying that the long hydrocarbon chains of DiI tightly embed into the nonpolar core of the SLB and limit its rotational diffusion. SMOLM imaging of another lipophilic dye, Merocyanine 540 (MC540), shows that it binds perpendicularly to a fluid (1,2-dioleoyl-sn-glycero-3-phosphocholine, DOPC) lipid membrane (Fig. 1b) with an orientation of θ=17.5±14.2° (Fig. 1d) and a narrower distribution of large solid angles (Ω=0.71π±0.22π sr), which agree with ensemble measurements.^36,37^ These observations confirm that SMOLM is capable of resolving both in-plane and out-of-plane molecules, as well as fixed and freely rotating dyes, within lipid membranes.

**Figure 1.**
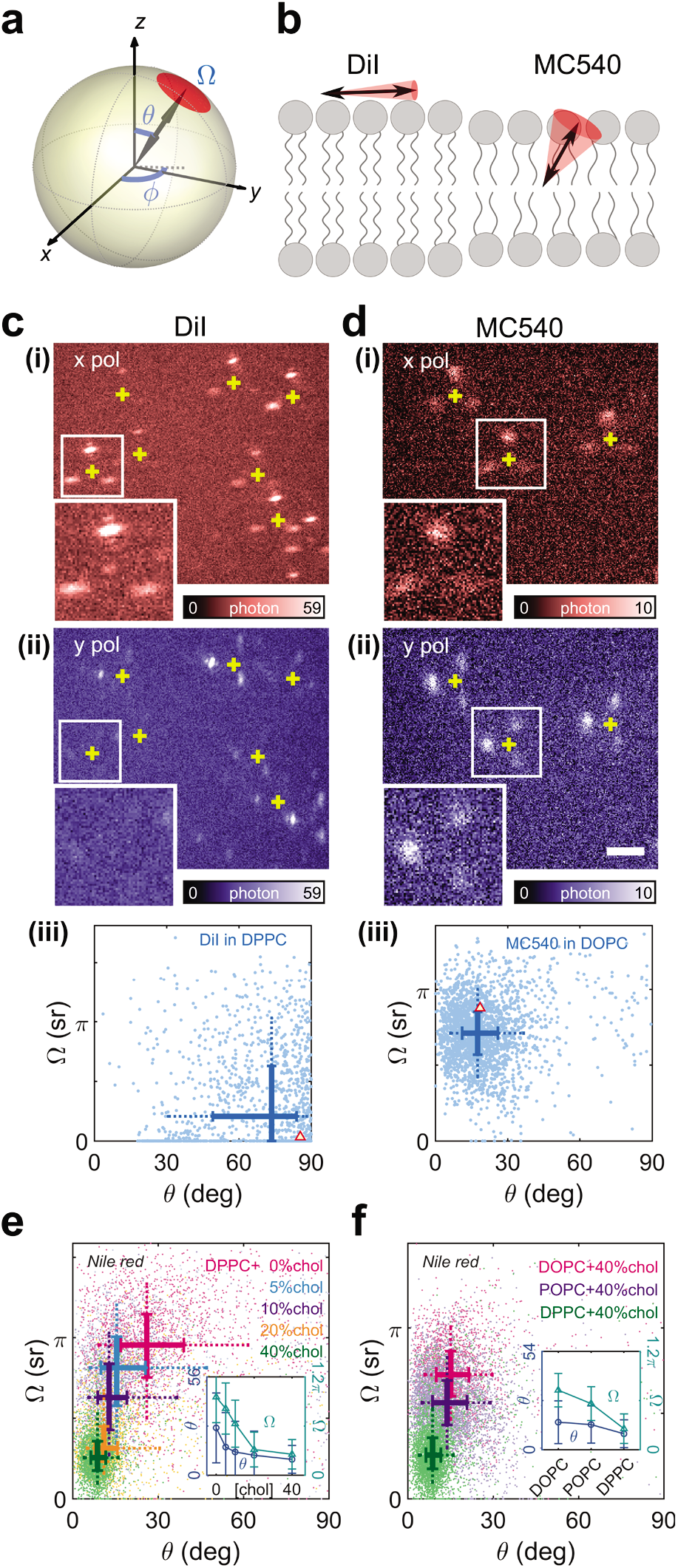
SMOLM imaging of the 3D orientations and wobbling of single fluorescent molecules. (**a**) Schematic of the 3D orientation and wobble of a single dipole, parameterized by polar angle (θ), azimuthal angle (ϕ), and wobble solid angle (Ω, modeling rotational diffusion within a cone). (**b**) Schematic of the orientation and wobble of DiI and MC540 within a supported lipid bilayer. (**c,d**) Representative images of DiI and MC540 in (**i**) x-polarized and (**ii**) y-polarized fluorescence channels using the Tri-spot point spread function (PSF). Yellow crosses represent the recovered position of each molecule. Insets: Magnified images of a single molecule. (**iii**) Orientation (polar angle θ) and wobble (solid angle Ω) measurements of single DiI and MC540 molecules. The thick solid lines in the scatter plots show the first to third quartile range of measured polar and solid angles; their intersection indicates the median values. The ends of dash lines indicate the 9th and 91st percentiles. The red triangle indicates the orientation spectra of the molecule shown in insets in **i** and **ii**. (**e,f**) Orientation (polar angle θ) and wobble (solid angle Ω) of Nile red in (**e**) DPPC with different cholesterol levels, and (**f**) DOPC, POPC, DPPC with various acyl chain structures. Insets: median polar angle and solid angle over different lipid conditions. Scale bar: 2 μm in **c,d**.

### Orientation spectra reveal lipid composition and packing

Within cell membranes, cholesterol (chol) plays a vital role in ordering and condensing lipid acyl chains, stabilizing lipid membranes, and forming nanoscale membrane domains.^1,38^ We discovered that the orientation spectra of single NR molecules are remarkably sensitive to the composition and packing of lipids influenced by chol. In DPPC without chol, NR exhibits a tilted out-of-plane orientation (θ=26.0±19.2°) and relatively large wobble (Ω=0.96π±0.31π sr). As the chol concentration increases to 40%, both polar and solid angles decrease drastically (θ=8.7±7.6°, Ω=0.26π±0.18π sr, Fig. 1e). These data suggest that chol strongly orders and condenses NR within the membrane in addition to the lipids themselves. Alternatively, we applied cholesterol-loaded methyl-β-cyclodextrin (10~400 μM, Supplementary Note 1) to elevate chol concentration *in-situ* within DPPC SLBs. The tilt and wobble of NR decrease to a level (Fig. S1a,b) commensurate with 40% chol. We observed the opposite effects on the orientation spectra of NR by adding melatonin (4%~30%, Supplementary Note 1, Fig. S1c), which is known to increase the disorder of lipid acyl chains and alleviate cholesterol’s effects.^39^

Our observations of NR’s orientational dynamics are remarkably consistent with the “umbrella model” of a lipid bilayer. In this model, the large hydrophilic phosphocholine headgroups form a cover, shielding cholesterol’s hydrocarbon steroid rings from the surrounding solvent while its hydroxyl group lies in close proximity to the lipid-water interface (Supplementary Note 2.1).^40^ Chol tends to align and condense lipid acyl chains, thereby restricting translational and rotational movements of molecules within the bilayer. Based on our observations, we surmise that NR primarily resides in the non-polar region of the bilayer surrounded by acyl chains. Chol-induced ordering orients NR parallel to neighboring acyl chains and perpendicular to the plane of the bilayer, and chol-induced condensation crowds molecules within the bilayer and thus decreases NR wobbling.

Interestingly, SMOLM reveals that the orientation dynamics of NR are more sensitive to the identity of lipid acyl chains than headgroups. In the presence of 40% chol with increasingly unsaturated lipids (DPPC, POPC (1-palmitoyl-2-oleoyl-glycero-3-phosphocholine) and DOPC, Fig. 1f), NR shows the largest solid (Ω=0.73π±0.21π sr) and polar (θ=15.1±11.8°) angles in disordered DOPC, compared to Ω=0.26π±0.18π sr and θ=8.7±7.6° in ordered DPPC. The disordered acyl chains likely counter chol’s ordering effect, thereby increasing the solid and polar angles of embedded fluorescent probes (Fig. S2a,b). In contrast, palmitoyl sphingomyelin (SPM) has the same acyl chains as DPPC but different headgroups. Interestingly, the orientation spectra of NR within SPM and DPPC are virtually indistinguishable with increasing chol concentration (Fig. 1e, S2c).

These SMOLM observations provide powerful insight into fluorophore interactions with lipid structures; in contrast to its structural analog Nile blue (Supplementary Note 2.2); NR emits fluorescence while inhabiting the non-polar region of a lipid bilayer, and its rotational dynamics are dictated by the specific environment “underneath the umbrellas”. Thus, SMOLM imaging reveals a molecule’s precise spatial positioning (<1 nm) within the lipid membrane, i.e., near headgroups vs. acyl chains, in addition to measuring the chemical environment surrounding each SM. Conventional SMLM does not have sufficient spatial resolution nor chemical sensitivity to visualize these properties.

### SMOLM imaging resolves lipid domains

Lipophilic probes that are sensitive to lipid packing enable SMOLM to map compositional and structural heterogeneities within lipid membranes, such as lipid domains. We carried out SMOLM imaging on a lipid mixture of DOPC/DPPC/chol. This mixture forms liquid-ordered (Lo) and liquid-disordered (Ld) phases as shown in conventional PAINT SMLM, where Lo/Ld domains are revealed by densities of MC540 localizations (Fig. 2a(i), dark: Lo, bright: Ld, Supplementary Note 3).^41^ SMOLM imaging, on the other hand, captures sensitive maps of chol concentration and acyl chain structure using the orientation spectra of NR (Fig. 2a(ii)(iii)). The Lo phases consist of ordered DPPC with chol,^1,42^ resulting in small NR solid and polar angles, compared to Ld phases formed by disordered DOPC (Fig. 2c). Since both orientation and wobble carry information useful for resolving Lo/Ld domains, we use principal component analysis (PCA, Experimental section) to combine the polar and solid angle data into a scalar “phase-index” map that discriminates Lo and Ld domains (Fig. 2a(iv), −0.28 arb. unit threshold).

**Figure 2.**
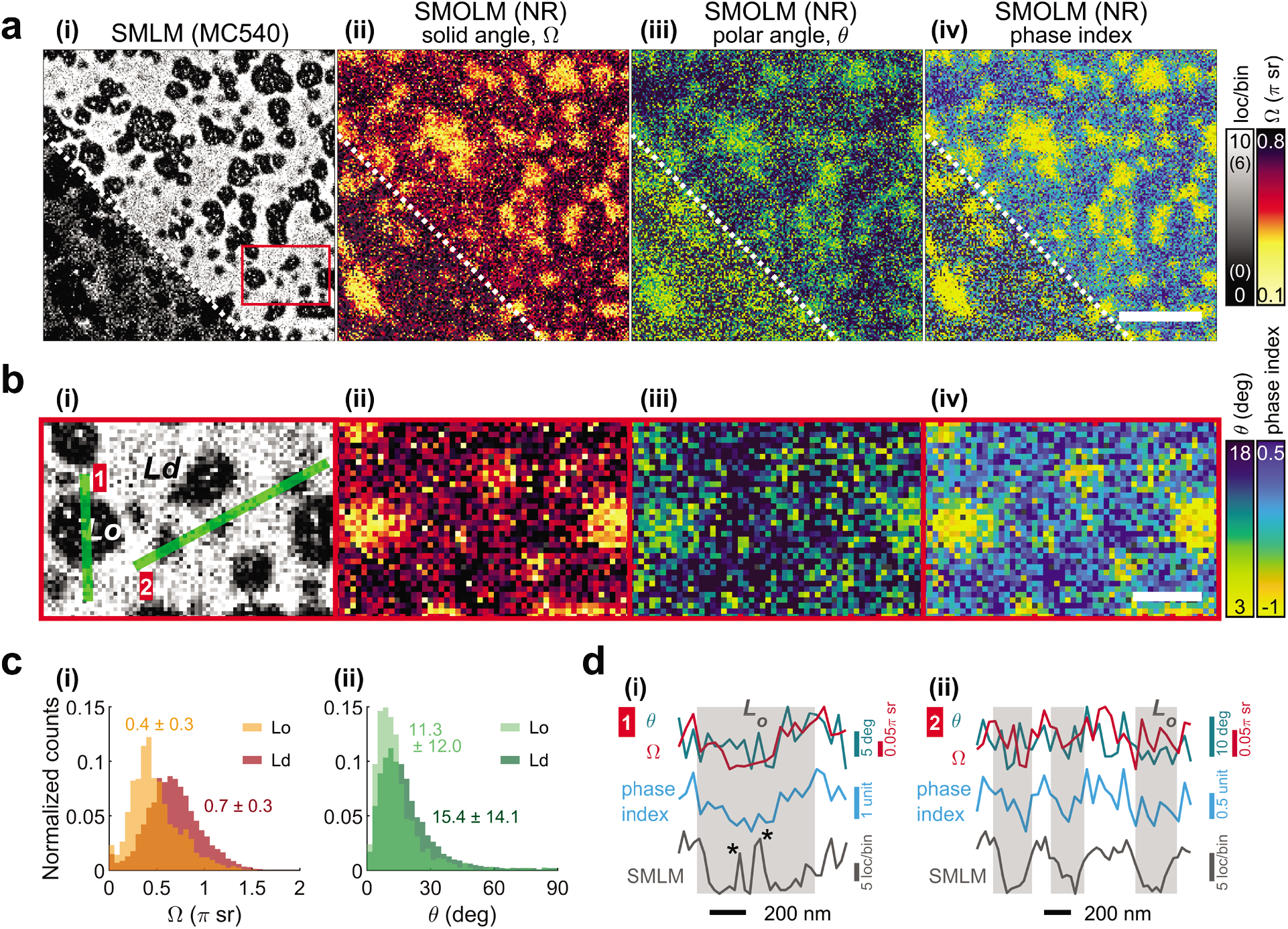
SMOLM imaging of Lo and Ld domains within a ternary lipid mixture of DOPC/DPPC/chol (35:35:30, molar ratio). (**a**) (**i**) Conventional MC540 SMLM image and (**ii-iv**) Nile red (NR) SMOLM images depicting (**ii**) solid angle (Ω), (**iii**) polar angle (θ), and (**iv**) combined phase index of a ternary lipid mixture of DOPC/DPPC/chol. Inset, lower-left: SMLM (colormap limits shown in parentheses) and SMOLM images filtered to show identical SM localization densities of 900 molecules/μm^2^. (**b**) Magnified view of (**i**) SMLM and (**ii-iv**) SMOLM images from boxed region in **a**. (**c**) Histogram and median±std of (**i**) solid angles and (**ii**) polar angles of Nile red in Lo and Ld domains in **b**. Gray pixels in SMOLM images represent bins with zero localizations. (**d**) Cross-sectional profiles of SMLM localizations, solid angle (Ω), polar angle (θ), and phase index along the green lines (1,2) in **b**. Gray shaded regions represent Lo domains. Scale bar: 2 μm in **a**, 500 nm in **b**. Bin size: 28 nm in SMLM, 40 nm in SMOLM.

SMOLM imaging shows Lo domains of various sizes both above (~500 nm) and below (<200 nm) the diffraction limit (green lines in Fig. 2b(i)). In large Lo domains, the cross-sectional profile of phase index (small values) matches well with the SMLM data (few localizations, Fig. 2d(i)). However, SMLM imaging is susceptible to contrast fluctuations from stochastic probe binding times, as shown by the local spikes (bins with dramatically more localizations) within Lo domains (Fig. 2d(i), marked by asterisks). These spikes are generated by MC540 with long binding times and confined lateral diffusion in Lo domains (Movie S1). On the other hand, SMOLM measures the orientation spectra of every probe molecule independent of the probe’s binding time to the membrane and is more robust to nonuniform localization densities over different lipid phases. SMOLM also shows good performance for resolving Lo domains below the diffraction limit. The cross-sectional profile of phase index within these small Lo domains is nosier but still consistent with the SMLM profile (Fig. 2d(ii)).

One advantage of SMOLM is that lipid composition and packing information are inferred from orientation measurements, which are collected simultaneously with molecule positions. If the position and orientation of each molecule are accurately estimated, only one measurement is required to distinguish Lo and Ld phase in a given pixel. Therefore, for a given SM localization density, SMOLM maps contain better contrast between Lo and Ld domains than SMLM images (Fig. 2a inset, Supplementary Note 4, Fig. S4).

To achieve optimal SMOLM imaging with high spatiotemporal resolution, one must select the best combination of orientation-sensitive probes and PSFs. First, we choose the probe whose orientation spectra are most separable between various single lipid phases. Between DOPC and DPPC, MC540 shows a larger separation in polar angle (ΔΘ_DPPC-DOPC_=55.6°, Fig. S5b) than that of NR (ΔΘ_DPPC-DOPC_=16.4°, Fig. S5f). Therefore, MC540 better discriminates gel versus liquid domains (Supplementary Note 5.1) in SMOLM. However, in the presence of chol, NR has superior performance to MC540 in distinguishing Lo and Ld domains (Supplementary Note 5.2). Next, one must choose a PSF that balances signal-to-background ratio (SBR), and therefore SM detection, with orientation sensitivity, i.e., the ability to resolve various orientational motions unambiguously.^27^ Due to varying measurement sensitivities, low SBRs, and tuning of analysis algorithms, different PSFs may perceive identical orientation spectra differently. However, these effects may be mitigated via instrument calibration (Supplementary Notes 6-8).

### Imaging enzyme-mediated changes to lipid composition

We next extend SMOLM to monitor *in situ* enzyme-mediated lipid compositional dynamics. In the plasma membrane, the hydrolysis of SPM via sphingomyelinase (SMase) generates a bioactive lipid, ceramide (cer), which selectively displaces chol from Lo domains at a 1:1 molar ratio,^43^ promotes lipid phase reorganization, forms a ceramide-rich ordered phase,^44^ and impacts cellular signaling and other vital processes.^45^ Most of these nanoscopic structural details were first observed by atomic force microscopy (AFM),^44^ which however is mostly limited to planar and static lipid samples and often requires complementary fluorescence imaging for visualizing lipid dynamics on faster timescales.^46,47^

Conventional SMLM imaging shows that SMase causes extensive changes in the morphology of ordered domains in DOPC/SPM/chol (Supplementary Note 9). However, for low SMase concentrations, the morphology of the Lo domains are mostly conserved and limited information on enzyme activity can be obtained. We instead applied SMOLM to monitor the underlying lipid compositional changes and resolve the spatial redistribution of newly generated ceramide and displaced chol within individual Lo domains.

A new orientation-sensitive Duo-spot PSF was developed for improved SBR over the Tri-spot PSF (Supplementary Note 6). We confirmed that NR SMOLM imaging using the Duo-spot PSF has excellent sensitivity for distinguishing newly generated ceramide domains vs. chol-rich Lo domains in static single-phase lipid samples (Supplementary Note 10.2). We next conducted SMOLM imaging of mixed DOPC/SPM/chol bilayers with successive SMase treatments of increasing dosage. Low SMase doses were chosen to test SMOLM sensitivity for detecting subtle enzyme activity within Lo domains (Fig. 3a, SMLM images at t_0_,t_3_). The SMOLM maps (Fig. 3b) indicate a dose-dependent disappearance of chol-rich Lo domains. Before treatment (t_0_), the shapes and positions of chol-rich Lo domains (small polar angle, solid angle, and phase index) imaged by SMOLM match the Lo domains mapped by SMLM. SMase treatment (16 mU/mL) induced insignificant changes in the SMOLM maps (Fig. 3b, t_1_), while more regions within the Lo domains begin to lose their chol-rich signature at a larger dose (50 mU/mL SMase, t_2_). After a 250 mU/mL dose, almost all the chol-rich Lo domains disappeared (Fig. 3b, t_3_); however, SMLM only reveals very minor changes in the size and shape of Lo domains (Fig. 3a, t_3_). The changes in orientation spectra agree well with those of NR within SPM+chol and SPM+cer lipid samples and strongly indicate the generation of cer-rich, chol-poor Lo domains (Fig. S13b, Table S2).^43^

**Figure 3.**
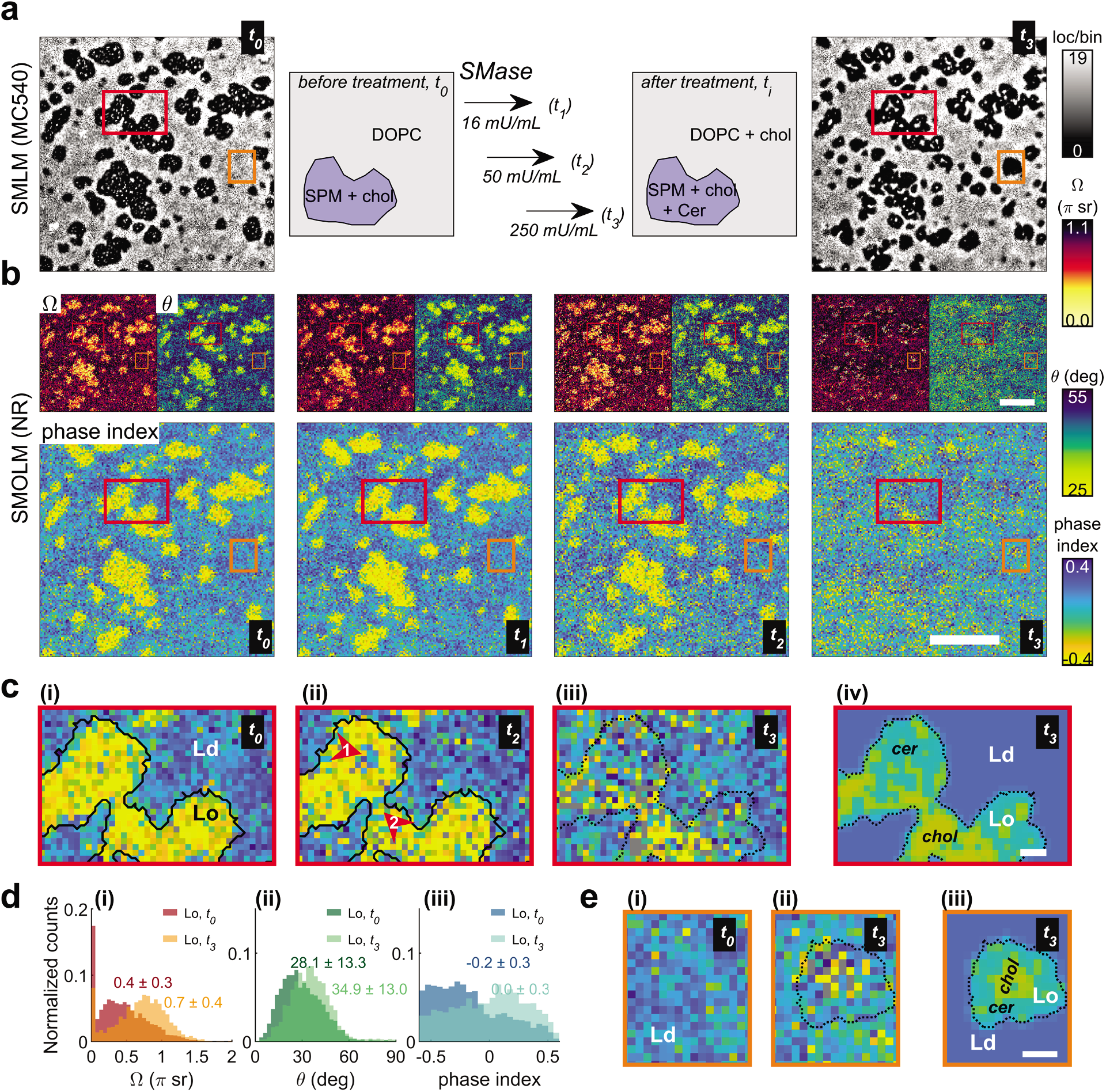
SMOLM imaging of SMase-induced lipid composition alteration and domain reorganization in a ternary lipid mixture DOPC/SPM/chol (35:35:30, molar ratio). (**a**) Conventional MC540 SMLM image and schematic of lipid mixture of DOPC/SPM/chol before (t_0_) and after (t_3_) three treatments of SMase. (**b**) SMOLM images (solid angle (Ω), polar angle (θ), and phase index) of Nile red before (t_0_) and after SMase treatments of 16 mU/mL (t_1_), 50 mU/mL (t_2_), and 250 mU/mL (t_3_). (**c**) Magnified view of the SMOLM phase-index map from the red boxed region in **a** and **b** at (**i**) t_0_, (**ii**) t_2_, and (**iii**) t_3_, and (**iv**) a lipid composition map at t_3_, indicating compositional changes within the Lo domain with minor changes in size and shape during SMase treatment. (**d**) Histogram and median±std of (**i**) solid angle, (**ii**) polar angle, and (**iii**) phase index of the single Lo domain in **c** before (t_0_) and after (t_3_) SMase treatment. (**e**) Magnified view of the SMOLM phase-index map from the orange boxed region in **a** and **b** at (**i**) t_0_ and (**ii**) t_3_ and (**iii**) a lipid composition map at t_3_, indicating a newly generated Lo domain by SMase. The solid and dotted black lines in **c**,**e** represent the Lo domain boundary before and after SMase treatment, repectively. Scale bar: 2 μm in **a**,**b**, 200 nm in **c**,**e**. Bin size: 28 nm in SMLM, 45 nm in SMOLM.

We focus our analysis on one particular Lo domain (red box in Fig. 3a,b). The boundary of the Lo domain before (solid black line in Fig. 3c) and after (dotted black line in Fig. 3c) SMase treatment was determined by the SMLM images. At a dose of 50 mU/mL (t_2_), the phase indices in several regions increase (arrows in Fig. 3c(ii)), indicating that these Lo regions are beginning to lose chol. This process is always localized in the interior of the domain (arrow 1, Fig. 3c(ii)) or at the intersection of two domains (arrow 2, Fig. 3c(ii)), which agrees with previous nanoscopic AFM observations.^44,46^ As the dose of SMase increases to 250 mU/mL (Fig. 3c(iii)), the chol-rich phase continues to shrink and is replaced by ceramide-rich domains. This compositional transition is also clearly illustrated by the change in orientation spectra from t_0_ to t_3_ (Fig. 3d).

To clearly visualize the lipid composition distribution, we designate the region outside of the domain boundary as the Ld phase and use the phase index threshold of −0.014 (arb. Units, Supplementary Note 10.2) to separate the chol-rich phase from the ceramide-rich phase (Fig. 3c(iv)). We also identified newly formed Lo domains (orange box in Fig. 3a,b), which are composed of well-separated chol-rich and ceramide-rich phases (Fig. 3e(ii)(iii)). The lipid composition maps among different Lo domains (Fig. 3c, 3e, Supplementary Note 12) reveal spatially heterogeneous nanoscale SMase activity, and also suggest that both SMase-generated ceramide and ceramide-displaced cholesterol rapidly (~seconds-minutes) condense into respectively enriched Lo domains.

## Discussion

A defining feature of soft matter is the impact of thermal fluctuations on the organization and self-assembly of molecules into mesoscopic structures like lipid membranes – processes that are notoriously difficult to observe directly. SMOLM extends conventional SMLM to measure both the positions and 3D orientations of single fluorescent molecules with high precision and sampling density (>3000 mol./μm^2^). We have utilized the orientation and rotational dynamics of fluorescent probes to reveal their interactions with the surrounding environment, such as the ordering of and condensation dynamics within lipid membranes. Our study demonstrates the feasibility of a new type of nanoscale imaging spectroscopy, namely measuring single-molecule orientation spectra, i.e., the six orientational second moments^27,49^ of dipole emitters, to resolve nanoscale chemical properties, similar to classic spectroscopies such as absorption, fluorescence emission, fluorescence lifetime,^50^ and NMR.

It has long been observed, using fluorescence polarization imaging of giant vesicles, that NR or Laurdan derivatives exhibit preferentially perpendicular orientations relative to the membrane surface in Lo phases due to constrained lipid packing and no preferential orientation in loosely packed Ld phases.^20–22^ Our SMOLM images provide the first quantitative measurements of this phenomena at the SM level and confirm that both polar angle and wobbling are increased in the Ld phase (Table S3). Leveraging this effect, SMOLM images (i.e., phase-index maps) can be used to discriminate between types of lipid domains (Supplementary Note 10). Compared to SMLM, SMOLM requires fewer total localizations and is less affected by localization density fluctuations when used for classifying Lo/Ld domains.

SMOLM relies on optimized orientation-sensitive PSFs to precisely measure orientation spectra and discover structural and chemical details of the sample. Fundamentally, to measure orientation with high sensitivity, the photons from each SM must be spread across multiple snapshots or camera pixels, thereby lowering the SBR compared to SMLM. Furthermore, rotational motions of fluorescent molecules are often accompanied by translational motions (diffusion), all of which are critical parameters to disentangle when probing molecular interactions in complex soft matter systems. Therefore, designing compact PSFs that can discriminate between translational and rotational diffusion, combined with new image analysis algorithms based upon machine learning,^48^ could further improve SMOLM’s spatiotemporal resolution for capturing faster biological processes. We anticipate that SMOLM will enable high throughput studies of both translational and orientational dynamics of single fluorescent probes within various soft matter systems, facilitate the discovery of mechanisms that control the orientation of individual molecules, and promote the design of new probes whose orientation conveys improved sensitivity and specificity for sensing various biophysical and biochemical phenomena.

## Experimental section

### Materials

1,2-dipalmitoyl-sn-glycero-3-phosphocholine (DPPC, 850355), 1,2-dioleoyl-sn-glycero-3-phosphocholine (DOPC, 850375), 1-palmitoyl-2-oleoyl-glycero-3-phosphocholine (POPC, 850457), *N*-palmitoyl-D-erythro-sphingosine (ceramide, 860516) were purchased from Avanti Polar Lipids. Cholesterol (C8667), cholesterol-loaded methyl-β-cyclodextrin (MβCD-chol, C4951), melatonin (M5250), *N*-palmitoyl-D-sphingomyelin (SPM, 91553), sphingomyelinase (S7651), Nile Blue (370088), glucose (G8270), glucose oxidase (G2133), catalase (C100), sodium chloride (S9625), Trizma base (T1503), hydrochloric acid (320331) were purchased from Sigma-Aldrich. Merocyanine 540 (M24571), Nile Red (AC415711000), DiI-C18(5) (D3911), carboxylate-modified microspheres (0.1 μm, 350/440 blue fluorescent, F8797) were purchased from Thermo Fisher Scientific. Deionized water (>18 MΩ·cm) was obtained through a Milli-Q water purification system and used for all aqueous solutions. High precision cover glass (No. 1.5H, thickness 170 μm±5 μm, 22×22 mm, Marienfeld) was used for all imaging.

### Imaging buffers

Tris buffer: 100 mM NaCl, 10 mM Tris, pH 7.4. GLOX buffer: 50 mM Tris (pH 8.3), 10 mM NaCl, 10% (w/v) glucose, and 1 % (v/v) enzymatic oxygen scavenger system. The stock solution of enzymatic oxygen scavenger system was prepared by adding 8 mg glucose oxidase and 38 μL 21 mg/mL catalase into 160 μL PBS, followed by 1 min 15,000 rpm centrifuging. The precipitate was removed before use.

### Supported lipid bilayer (SLB) preparation

Supported lipid bilayers (SLB) were formed by fusing vesicles on coverslips. To prepare large unilamellar vesicles (LUVs), a lipid mixture was first dissolved in chloroform, followed by evaporation of the solvent and drying for over 12 h under vacuum. The lipids were resuspended by adding Tris buffer (100 mM NaCl, 3 mM Ca^2+^, 10 mM Tris, pH 7.4) to arrive at a final lipid concentration of 1 mM. The lipid suspension was vigorously vortexed for 30 min under nitrogen before extrusion (25 passages, Avanti Polar Lipids). The monodisperse LUVs were next added onto ozone-cleaned (UV Ozone Cleaner, Novascan Technologies) coverslips and incubated in a water bath at a temperature higher than the phase transition temperature of the lipids for 1 hour to form a SLB. After 30 min of cooling to room temperature, the lipid bilayer was thoroughly rinsed with Tris buffer to remove residual lipids and imaged immediately. To track the possible shift of coverslip during super-resolution imaging, a layer of fluorescent beads (0.1 μm, blue fluorescent, 1:2000 dilution in H_2_O) were sparsely spin-coated (2500 rpm for 40 s) on the coverslip before the deposition of the SLB.

### SMOLM imaging

A home-built microscope^27^ with a 100× objective lens (NA 1.40, Olympus, UPLSAPO100XOPSF) was used to perform SMOLM imaging. For NR and MC540 imaging, a 561-nm laser (Coherent Sapphire) with a peak intensity of 1.31 kW/cm^2^ and a dichroic beamsplitter (Semrock, Di03-R488/561) were used. The emission was filtered by a bandpass filter (Semrock, FF01-523/610), and separated into x- and y-polarized channels by a polarization beam splitter (PBS, Meadowlark Optics, BB-100-VIS). The Tri-spot and Duo-spot phase masks were generated by a spatial light modulator (Meadowlark Optics, 256 XY Phase Series) onto which the back focal plane of both polarization channels was projected. The modulated SMOLM images captured with a typical 30 ms integration time using an sCMOS camera (Hamamatsu ORCA-flash4.0 C11440-22CU). A 514-nm laser (Coherent Sapphire) with peak intensity of 1.56 kW/cm^2^, dichroic beamsplitter of Di02-R514, and bandpass filter of FF01-582/64 were used for DiI imaging. Tris buffer or GLOX buffer (only for MC540) was used as imaging buffer.

### Data analysis

The raw images were analyzed by using a bespoke maximum-likelihood estimation algorithm with joint-sparse regularization^28,29^ written in Matlab. The position, brightness, orientation (polar angle, azimuthal angle) and wobble (solid angle) of each fluorescent molecule were estimated simultaneously. Since a finite number of signal photons leads to a bias in the largest detectable solid angle, the plotted range of each figure showing measured wobble angles was adjusted based on the measured signal and background photons in each particular experiment.^51^ Principal component analysis (PCA) and kernel principal component analysis (KPCA) (scripts written in Python and available online^52^) were used to combine the polar and solid angle data. The PCA scores quantify the separation of different lipid phases (Supplementary Note 10) and were thus designated as “phase indices”. To generate SMOLM maps, the localizations were first grouped with appropriate bin size. In each bin, the median values (unless otherwise stated, see Supplementary Note 11) of polar angle, solid angle, and phase index were calculated and assigned to each bin.

## Acknowledgements

Research reported in this publication was supported by the National Science Foundation under grant number ECCS-1653777 and by the National Institute of General Medical Sciences of the National Institutes of Health under grant number R35GM124858. The authors acknowledge financial support from Washington University in St. Louis and the Institute of Materials Science and Engineering for the use of instruments and staff assistance. Computations were performed using the facilities of the Washington University Center for High Performance Computing, which were partially funded by NIH grants 1S10RR022984-01A1 and 1S10OD018091-01.

## Author contributions

J.L. and M.D.L. designed the research; J.L. designed and performed the experiments; J.L., H.M., T.D., and O.Z. implemented analysis algorithms and performed data analysis; T.D. designed and characterized the Duo-spot PSF; M.D.L. supervised the research; J.L., T.D., and M.D.L. wrote the manuscript; all coauthors discussed and commented on the manuscript.

