## Supplementary Information for "Single-molecule 3D orientation imaging reveals nanoscale compositional heterogeneity in lipid membranes"

**Supplementary Note 1. Orientation spectra of Nile red in various lipid environments** 3

**1.1 Supported lipid bilayers (SLBs) containing cholesterol (chol) and melatonin (mel)** 3

**1.2 Lipids of various acyl chains and head groups** 4

**Supplementary Note 2. Sensitivity of Nile red orientation spectra to cholesterol concentration** 5

**2.1 Mechanistic insight provided by the "umbrella model"** 5

**2.2 SMOLM imaging of Nile blue within SLBs** 5

**Supplementary Note 3. SMLM imaging of Merocyanine 540 to resolve lipid phases** 7

**Supplementary Note 4. Impact of localization density on SMOLM and PAINT SMLM imaging for distinguishing Lo and Ld domains** 8

**Supplementary Note 5. Selecting probes for better lipid domain resolvability in SMOLM** 11

**5.1 Resolving gel and liquid domains** 11

**5.2 Resolving Lo and Ld domains** 14

**Supplementary Note 6. Selection of orientation-sensitive PSFs in SMOLM** 15

**6.1 Localization and orientation estimation of SMOLM** 15

**6.2 PSF detectability** 17

**6.3 Design of the Duo-spot PSF** 18

**6.4 SMOLM performance using the Duo-spot PSF for measuring chol in SLBs** 22

**Supplementary Note 7. Molecular lateral diffusion distorts single molecule PSFs and biases orientation and wobble estimation** 23

**Supplementary Note 8. Comparison of the Duo-spot vs. Tri-spot PSFs for SMOLM imaging of DOPC/SPM/chol SLBs** 26

**Supplementary Note 9. SMLM of SMase-induced lipid domain reorganization** 30

**Supplementary Note 10. Principal component analysis (PCA) analysis of SMOLM data** 31

**10.1 PCA analysis to distinguish gel (or Lo) and liquid (or Ld) phases** 31

**10.2 KPCA analysis to distinguish lipid phases in DOPC/SPM/chol with SMase treatment** 31

**Supplementary Note 11. SMOLM data visualization using the Duo-spot PSF** 35

**Supplementary Note 12. SMase-mediated lipid composition changes in Lo domains** 37

**Supplementary Note 13. Orientation spectra statistics of single-phase SLBs** 39

**Supplementary Movies** 42

**References** 43

### Supplementary Note 1. Orientation spectra of Nile red in various lipid environments

#### 1.1 Supported lipid bilayers (SLBs) containing cholesterol (chol) and melatonin (mel)

The concentration of cholesterol (chol) in DPPC SLBs was elevated *in-situ* using cholesterol-loaded methyl-β-cyclodextrin (MβCD-chol) (dissolved in Tris buffer). After four successive MβCD-chol treatments, the out-of-plane tilt and wobble of Nile red (NR) decrease to a level (θ=8.4±8.2°, Ω=0.20π±0.14π sr, Fig. S1a) commensurate with 40% chol (Fig. 1e). Repeating treatments with high doses of MβCD-chol has little impact on the orientation spectra, implying that the solubility limit of chol has been reached (Fig. S1b).

Different amounts of melatonin (mel) were added to the lipid mixture during the preparation of lipid SLBs. Adding melatonin results has an opposite effect on the orientation spectra of Nile red (Fig. S1c); both the tilt and wobble angles increase.

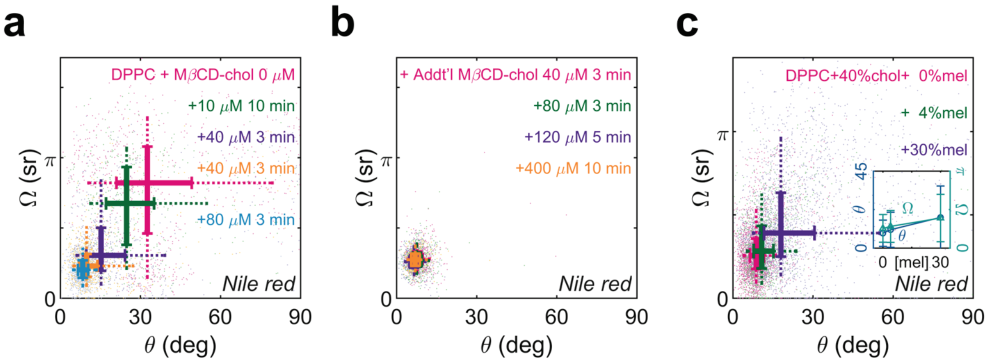


Fig. S1 | (**a**) Orientation (polar angle θ) and wobble (solid angle Ω) of Nile red in DPPC after the treatment of MβCD-chol with different dose and duration. (**b**) Orientation and wobble of Nile red after additional treatment of higher dose of MβCD-chol, implying the maximum ordering effect that chol can impart upon Nile red, which most likely corresponds to the maximum equilibrium solubility of cholesterol in DPPC. (**c**) Orientation and wobble of Nile red in DPPC+40% chol with different levels of melatonin. Insets: changes in median polar angle and solid angle for various melatonin concentrations. The thick solid lines in the scatter plots show the first to third quartile range of measured polar and solid angles; their intersection indicates the median values. The ends of dash lines indicate the $9^{th}$ and $91^{st}$ percentiles.

#### 1.2 Lipids of various acyl chains and head groups

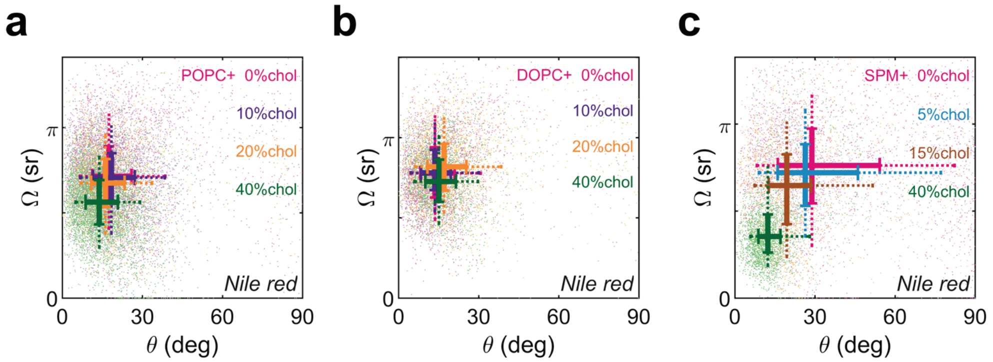


Fig. S2 | Orientation (polar angle θ) and wobble (solid angle Ω) of Nile red in (**a**) POPC, (**b**) DOPC, and (**c**) SPM SLBs with various concentrations of cholesterol.

### Supplementary Note 2. Sensitivity of Nile red orientation spectra to cholesterol concentration

#### 2.1 Mechanistic insight provided by the "umbrella model"

In umbrella model of a lipid bilayer,[1] the large hydrophilic phosphocholine headgroups form a cover to shield the hydrocarbon steroid rings of cholesterol and prevent their exposure to water.[1] Therefore, chol is expected to be positioned in a configuration with its hydroxyl group in close proximity to the lipid-water interface and its hydrocarbon steroid incorporated into the nonpolar interior of the lipid membrane. When Nile red binds to the lipid membrane, it is reported to also occupy the interfacial region of the membrane as chol.[2] The ordering effect of chol on the lipids within the bilayer, as well as the noncovalent interactions between the planar 4-ring structure of chol and planar benzophenoxazine of Nile red, causes Nile red to align along the orientation of chol. The "alignment" effect is so strong that even with a very small amount of chol (e.g. 5%) present, a dramatic decrease in polar angle of Nile red is observed (Fig. 1e inset, $\theta$ curve). Moreover, as chol concentration increases, chol takes more space in the hydrophobic regions under the phospholipid polar headgroup, which reduces the free volume and limits the wobbling movement of Nile red. This "crowding" effect is proportional to the amount of chol added to the lipid membrane, which explains the inverse relationship between solid angle and chol concentration (Fig. 1e inset, $\Omega$ curve).

#### 2.2 SMOLM imaging of Nile blue within SLBs

Nile blue is an analogue of Nile red in the benzophenoxazine family (Fig. S3a),[3] and as a cationic dye, it is more likely than Nile red to stay in the polar head region of the lipid bilayer and avoid entering the hydrophobic interior. We observed less changes in the orientation spectra of Nile blue ($\theta<5°$ and $\Omega<0.08\pi$ sr) in response to various lipid compositions and degrees of packing (Fig. S3b,c) compared to Nile red, which implies that Nile blue is insensitive to chol-induced ordering and condensation of the hydrophobic core of the bilayer. The observation is consistent with our aforementioned mechanistic view of the lipid "umbrella".

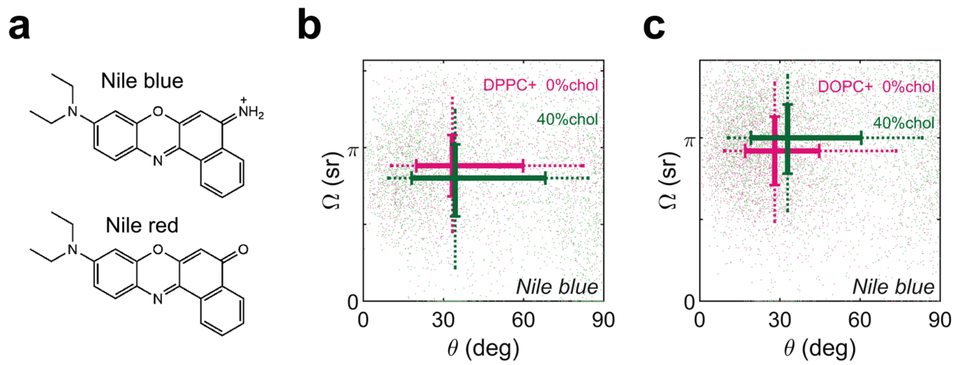


Fig. S3 | (**a**) Chemical structures of Nile blue and Nile red. (**b,c**) Orientation (polar angle θ) and wobble (solid angle Ω) of Nile blue in (**b**) DPPC with 0 and 40% chol and (**c**) DOPC with 0 and 40% chol.

### Supplementary Note 3. SMLM imaging of Merocyanine 540 to resolve lipid phases

Using points accumulation for imaging in nanoscale topography (PAINT, a form of conventional SMLM), Merocyanine 540 (MC540) is capable of resolving gel (or liquid-ordered, Lo) and liquid (or liquid-disordered, Ld) phases in lipid membrane[4]. Previous studies have found both fluorescent monomers and nonfluorescent dimers of MC540 in lipid membranes. Further, the monomer-dimer equilibrium is sensitive to lipid phase, where the equilibrium dimerization constant for MC540 in the liquid phase ($K_d=4\times10^3$ $M^{-1}$) is smaller than that in the gel phase ($K_d=1.7\times10^5$ $M^{-1}$).[5–7] Therefore, in SMLM, the number density of MC540 in gel (or Lo) phases is lower than liquid (or Ld) phases.[4] Typically, we captured 50,000 MC540 SMLM frames at an average localization density of 0.14 $\mu m^{-2}$ in a GLOX buffer (50 mM Tris, 10 mM NaCl, 10% w/v glucose, 0.5 mg/mL glucose oxidase, 0.05 mg/mL catalase, pH 8.0) used to minimize photobleaching. (Note that we found GLOX ineffective for reducing Nile red photobleaching, however.) Regions with dramatically fewer localizations are identified as gel (or Lo) domains, and brighter regions are labeled as liquid (or Ld) domains (Fig. 2a(i), Fig. 3a); note, however, that the localization threshold for determining the precise identity of these domains has not yet been characterized as a function of buffer, photoexcitation, and fluorescence acquisition conditions. In this work, we only use MC540 for SMLM imaging for distinguishing gel (or Lo) and liquid (or Ld) phases in lipid membrane.

### Supplementary Note 4. Impact of localization density on SMOLM and PAINT SMLM imaging for distinguishing Lo and Ld domains

In SMOLM, as long as the position and orientation of each molecule are accurately estimated, only one molecule, or one orientation measurement, is required in each bin to distinguish Lo and Ld domains from one another. For a given localization density, SMOLM reconstructs lipid domains and resolves their structure more robustly compared to SMLM.

To demonstrate, we use the data from the lipid mixture of DOPC/DPPC/chol (35:35:30, molar ratio) containing both Lo and Ld domains in Figure 2 in the main text. We randomly selected different subsets of localizations from the entire dataset and reconstructed the SMLM images and SMOLM maps. Each subset contains localizations corresponding to densities of 50, 100, 400, and 900 molecules/$\mu m^2$ (mol./$\mu m^2$). To guarantee the estimated position and orientation of the selected localizations are accurate, we only choose the localizations with the brightness higher than the third quartile of the entire population, i.e., the best localizations in both SMLM and SMOLM. For SMOLM data, the median value of phase index from each selected subset is same as the entire population to avoid a biased selection of localizations from Lo or Ld phases.

The MC540 SMLM image (Fig. S4a) with high localization density ($9\times10^3$ mol./$\mu m^2$) was used as the ground truth image to show the location, size, and shape of Lo and Ld domains. For low localization density (i.e. 50 mol./$\mu m^2$), the SMLM image (Fig. S4b(i)) has very limited resolution to reveal Lo domains. The corresponding phase-index map (Fig. S4c(i)) in the SMOLM image, however, already shows the locations of major Lo domains. As the localization density increases (100 to 900 mol./$\mu m^2$), both the SMLM images (Fig. S4b(ii)-(iv)) and SMOLM maps (Fig. S4c(ii)-(iv)) show improved contrast between the Lo and Ld domains.

To quantify the performance of SMOLM and SMLM to resolve the Lo/Ld phases, we measure the image similarity between the SMOLM or SMLM images under various localization densities and the ground truth image. First, we created binary Lo domain maps from SMOLM or SMLM data to illustrate the Lo/Ld domains. Regions in the SMLM image with fewer than 1 localization per bin, in the SMOLM image with phase index smaller than -0.28 (arb. Units, see also Supplementary Note 10.1), and in the ground truth SMLM image with fewer than 10 localizations per bin were designated as the Lo phase (blue regions in insets of Fig. S4a,b,c). Next, for each scenario with a given localization density (50 to 900 mol./μm$^2$), we calculated the Root Mean Squared Error (RMSE) between 1) the binary Lo domain map from the ground truth image (inset of Fig. S4a) and 2) the binary Lo domain map from SMOLM (insets of Fig. S4c) or SMLM (insets of Fig. S4b). As shown in Fig. S4d, $RMSE_{SMOLM}$ is smaller than $RMSE_{SMLM}$ below a localization density of 400 mol./μm$^2$. $RMSE_{SMLM}$ quickly drops as localization density increases and is lower than $RMSE_{SMOLM}$ at the localization density of 900 mol./μm$^2$. Note that since we designated the SMLM image as the ground-truth image in this analysis, we expect $RMSE_{SMLM}$ to outperform $RMSE_{SMOLM}$ at some threshold localization density, which is not an indicator of the true accuracy of discriminating Lo vs. Ld domains. In the future, the true RMSE performance can be determined by correlating against a secondary imaging modality.

Therefore, for a given total localization number, especially for low localization density, our data indicate that the SMOLM map exhibits better performance to distinguish Lo and Ld domains than conventional lipid membrane SMLM imaging via PAINT.

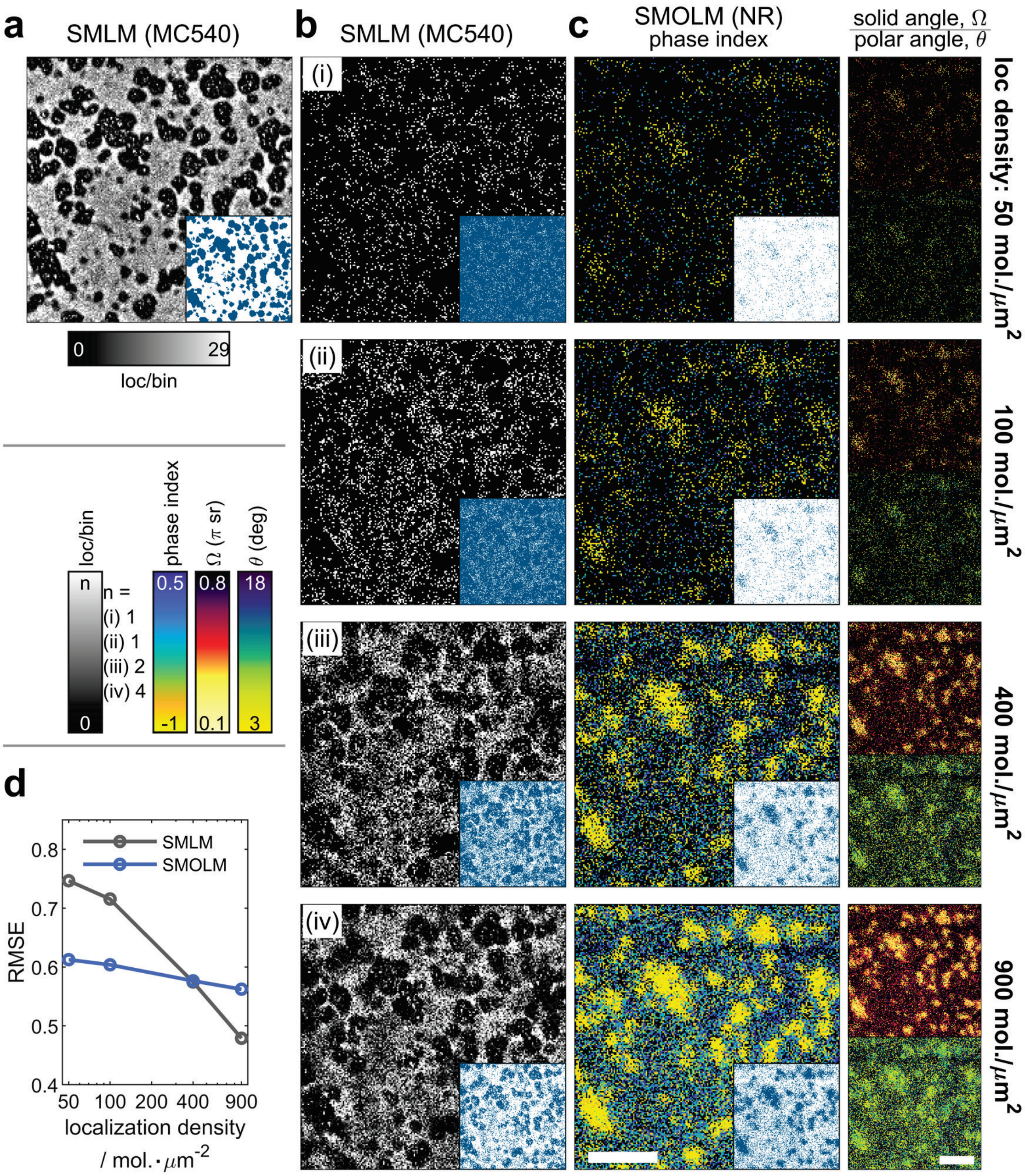


Fig. S4 | (**a**) Conventional MC540 SMLM image ($9\times10^3$ mol./μm$^2$ localization density) of a ternary lipid mixture of DOPC/DPPC/chol showing Lo (dark) and Ld (bright) domains. (**b**) MC540 SMLM and (**c**) Nile red SMOLM images reconstructed from subset of the raw data with various localization densities: (**i**) 50, (**ii**) 100, (**iii**) 400, and (**iv**) 900 mol./μm$^2$. Insets in **a**,**b**,**c**: Lo domain maps (blue, binary images) constructed from MC540 SMLM and Nile red SMOLM images with corresponding localization densities. (**d**) Root Mean Squared Error (RMSE) between the ground truth Lo domain map in inset of **a** and SMLM or SMOLM Lo domain maps in insets of **b** or **c**. Scale bar: 2 μm. Bin size: 45 nm in **a**,**b**,**c**.

### Supplementary Note 5. Selecting probes for better lipid domain resolvability in SMOLM

#### 5.1 Resolving gel and liquid domains

To achieve better SMOLM imaging, one must select the probe whose orientation spectra across various lipid phases are most separable. For example, to optimize SMOLM for resolving gel and liquid phases within a mixture of DOPC/DPPC (1:1, molar ratio), we measured the orientation spectra of MC540 and Nile red in a single-component DPPC (gel) SLB and a DOPC (liquid) SLB.

MC540 exhibits preferential orientations in different lipid membrane phases, which was previously measured for an ensemble or bulk collection of molecules.[5,6] We used SMOLM to measure the orientation of MC540 at the single-molecule (SM) level (Fig. S5b). In DPPC (gel phase), our results indicate that MC540 exhibits large polar angles ($\theta$=73.1±23.7°) and a wide distribution of solid angles ($\Omega$=0.68$\pi$±0.41$\pi$ sr); whereas in DOPC (liquid phase), MC540 shows small polar angles ($\theta$=17.5±14.2°) and a narrower distribution of solid angles ($\Omega$=0.71$\pi$±0.22$\pi$ sr). The results match well with the literature reports.[8]

The orientation of single Nile red molecules within lipid membranes has never been measured before. Our SMOLM results (Fig. S5f) indicate that Nile red exhibits out-of-plane orientations ($\theta$=15.1±11.3°) in DOPC (liquid phase) SLBs with a solid angle ($\Omega$) of 0.82$\pi$±0.24$\pi$ sr. The polar angle ($\theta$) of Nile red increases to 31.5±21.4° in DPPC (gel phase) SLBs with slightly deceased solid angles ($\Omega$=0.81$\pi$±0.38$\pi$ sr) (Fig. S5f).

Our measurements show that the polar angle of MC540 exhibits larger separation ($\Delta\theta_{DPPC\text{-}DOPC}$=55.6°) in DOPC versus DPPC than Nile red ($\Delta\theta_{DPPC\text{-}DOPC}$=16.4°). We therefore compared SMOLM imaging using MC540 (Fig. S5a-d) versus Nile red (Fig. S5e-h) for a mixed DOPC/DPPC SLB (1:1, molar ratio) from the same field of view.

In the mixture of DOPC/DPPC, conventional MC540 SMLM resolves both gel (dark regions in Fig. S5a(i)) and liquid (bright regions in Fig. S5a(i)) domains. As expected from the single-component lipid imaging (Fig. S5b), the gel domains within the SMOLM maps have larger polar angles and similar solid angles but a broader distribution (Fig. S5a(ii)(iii)) than liquid domains. The size and shape of resolved gel and liquid phases in the phase-index map (Fig. S5a(iv)) match those in the SMLM image (Fig. S5a(i)), which is further demonstrated by line profiles of selected Lo domains (green regions 1 and 2 in Fig. S5a(i)). The phase-index profiles are in good agreement with the profiles of SMLM image for both >1 μm (Fig. S5c(i)) and ~200 nm (Fig. S5c(ii)) Lo domains. We also plotted the histogram of solid angles (Fig. S5d(i)) and polar angles (Fig. S5d(ii)) from all localizations within the gel and liquid phases in region 1. The gel phase exhibits an additional population of MC540 with ~70° polar angle and a small population of zero sr solid angle, which matches the orientation spectra observed in single-component DPPC SLBs (Fig. S5b).

Following MC540 imaging, we gently washed the sample with buffer and performed SMOLM imaging using Nile red. The polar angles of Nile red within gel and liquid phases are less resolvable from one another than those of MC540, therefore producing a lower-quality phase-index map (Fig. S5e(iv)). Some gel domain features are missing compared to MC540 SMOLM images (Fig. S5a(iv)) and conventional SMLM images (Fig. S5e(i)). In addition, the phase-index line profile does not match the SMLM profile (blue and gray profiles in Fig. S5g), and the solid and polar angle histograms of gel and liquid phases (Fig. S5h) are indistinguishable.

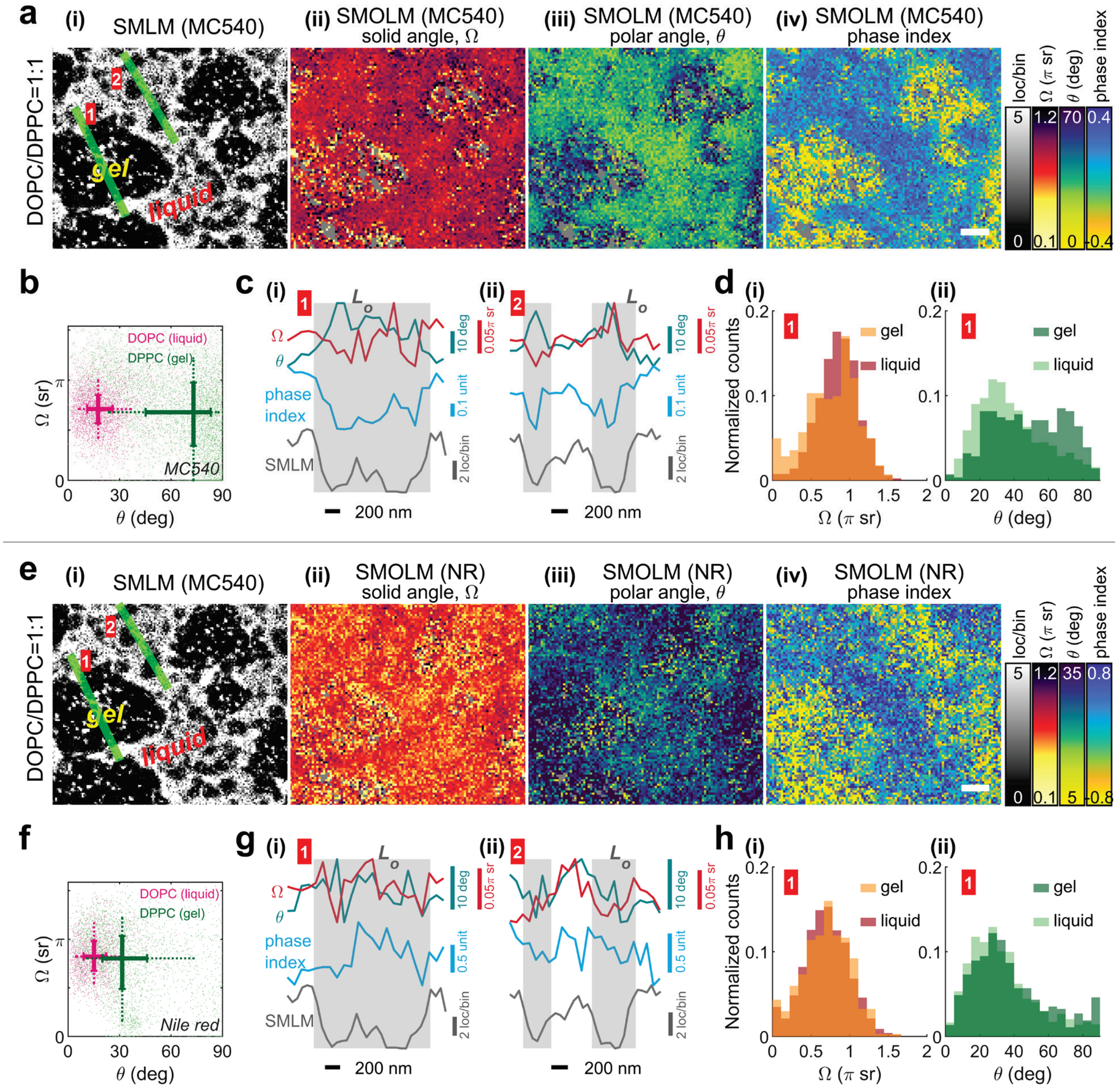

Fig. S5 | SMOLM imaging using MC540 and Nile red of a mixed DOPC/DPPC SLB (1:1, molar ratio). (**a**) (**i**) Conventional SMLM image and SMOLM images of (**ii**) solid angle (Ω), (**iii**) polar angle (θ), and (**iv**) phase index using MC540. (**b**) Single-phase lipid samples. Orientation (polar angle θ) and wobble (solid angle Ω) of MC540 in a DOPC (liquid phase) and a DPPC (gel phase) SLB. (**c**) Cross-sectional profiles of SMLM localizations, solid angle (Ω), polar angle (θ), and phase index along the green lines (**i**) **1** and (**ii**) **2** in **a**. Gray shaded regions represent Lo domains. (**d**) Histogram of (**i**) solid angle and (**ii**) polar angle in the region of green line **1**. (**e-h**) Corresponding SMOLM results of Nile red in the same field of view of the DOPC/DPPC mixture in **a-d**. Scale bar: 500 nm in **a**,**e**. Bin size: 28 nm in SMLM, 40 nm in SMOLM.

### 5.2 Resolving Lo and Ld domains

A second example is to characterize how MC540 and NR recognize Lo and Ld phases in a lipid mixture containing chol, such as DOPC/DPPC/chol. In such a mixture, chol is concentrated within DPPC domains to form the Lo phase, and DOPC forms the Ld phase. We measured the orientation spectra of Nile red and MC540 in a single-phase DPPC+chol (Lo) SLB and a DOPC (Ld) SLB (Fig S6). Chol greatly reduces the polar angle of MC540 in the Lo phase and makes it hardly distinguishable ($\Delta\theta_{\text{DPPC-DOPC}}$=2.3°, $\Delta\Omega_{\text{DPPC-DOPC}}$=0.11π sr) from that in the Ld phase (Fig, S6a, compared to Fig. S5b). Conversely, both the polar and solid angles of Nile red are well separated ($\Delta\theta_{\text{DPPC-DOPC}}$=4.9°, $\Delta\Omega_{\text{DPPC-DOPC}}$=0.52π sr) in Lo vs. Ld phases (Fig, S6b, compared to Fig. S5f). The data indicate Nile red is superior to MC540 in distinguishing Ld versus Lo domains. We therefore use Nile red for SMOLM imaging on DOPC/DPPC/chol and DOPC/SPM/chol lipid membranes as shown in Fig. 2 and Fig. 3 in the main text.

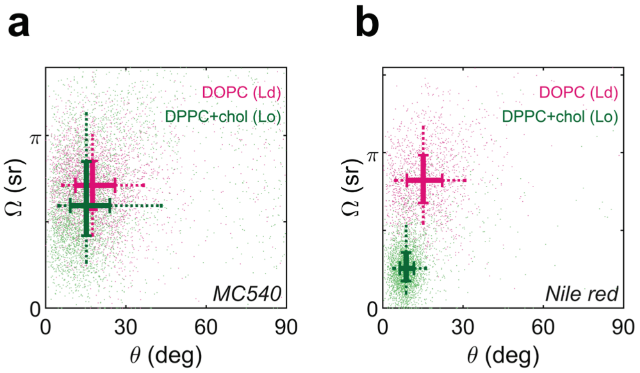


Fig. S6 | Orientation (polar angle θ) and wobble (solid angle Ω) of (**a**) MC540 and (**b**) Nile red in DOPC (Ld phase) and DPPC+chol (Lo phase) SLBs.

The observations and discussions above emphasize the importance of choosing fluorescence probes with the most separable orientation spectra in order to achieve well-resolved SMOLM imaging of lipid phase, composition, and/or packing.

### Supplementary Note 6. Selection of orientation-sensitive PSFs in SMOLM

#### 6.1 Localization and orientation estimation of SMOLM

We model a fluorescent molecule as a dipole-like emitter wobbling within a cone.[9,10] An orientational unit vector $\boldsymbol{\mu} = [\mu_x, \mu_y, \mu_z]^{\mathrm{T}} = [\sin\theta\cos\phi, \sin\theta\sin\phi, \cos\theta]^{\mathrm{T}}$ and solid angle $\Omega$ define the center orientation and the wobbling area of the cone, respectively (Fig. 1a). Assuming that a molecule's rotational correlation time is faster than its exited state lifetime and the camera acquisition time, its orientation state can be fully characterized by a second-moment vector $\boldsymbol{m} = [\langle\mu_x^2\rangle, \langle\mu_y^2\rangle, \langle\mu_z^2\rangle, \langle\mu_x\mu_y\rangle, \langle\mu_x\mu_z\rangle, \langle\mu_y\mu_z\rangle]^{\mathrm{T}}$, where each component is a time-averaged second moment of $\boldsymbol{\mu}$ within a single camera acquisition period.[11] A fluorescence microscope image $\boldsymbol{I} \in \mathbb{R}^n$ of such an emitter captured by an $n$-pixel camera can be modeled as a linear superposition of six basis images weighted by $\boldsymbol{m}$ as follows:

$$\boldsymbol{I} = s\boldsymbol{B}\boldsymbol{m} + b = s[\boldsymbol{B}_{xx}, \boldsymbol{B}_{yy}, \boldsymbol{B}_{zz}, \boldsymbol{B}_{xy}, \boldsymbol{B}_{xz}, \boldsymbol{B}_{yz}]\boldsymbol{m} + b, \tag{1}$$

where $s$ is the number of photons detected from the molecule and $b$ is the number of background photons in each pixel. Each so-called basis image $\boldsymbol{B}_k \in \mathbb{R}^n$ ($k \in \{xx, yy, zz, xy, xz, yz\}$) corresponds to the response of the optical system to each orientational second-moment component $m_k$ and can be calculated by the vectorial diffraction theory.[11–13]

In this paper, the locations and orientations of single molecules were estimated simultaneously using a sparsity-promoting maximum likelihood estimator.[14,15] Briefly, the object space is represented by a rectangular lattice of grid points with spacing equal to the camera pixel size (58.5 nm). Each grid point may contain at most a single molecule parameterized by brightness, position offsets, and six orientational second moments.

To robustly estimate these parameters at each grid point in the presence of SM image overlap, we use a regularized maximum likelihood exploiting a group-sparsity norm. The algorithm begins by estimating the strength, i.e., brightness, of each of the second moments $\tilde{m}_k$ independently at all object grid points. We next pool together localizations, i.e., their brightnesses and position offsets, across the six second moments to identify the most likely molecules in the object space. Once we identify these molecules, we solve a constrained maximum likelihood to minimize systematic biases induced by the sparsity norm, yielding estimates of the brightnesses, locations, and orientations (second moments $\widetilde{\boldsymbol{m}}$) of all molecules in the image. We remove localizations with signal estimates less than 400 photons detected to eliminate unreliable localizations.

The estimated second-moment vectors $\widetilde{\boldsymbol{m}}$ were next projected to an angular orientation space (polar angle $\theta$, azimuthal angle $\phi$, and wobbling area $\Omega$ of a transition dipole moment $\boldsymbol{\mu}$) by a weighted least-square estimator:

$$(\theta, \phi, \Omega) = \underset{\hat{\theta},\hat{\phi},\hat{\Omega}}{\operatorname{argmin}} \left(\widetilde{\boldsymbol{m}} - \boldsymbol{m}(\hat{\theta}, \hat{\phi}, \hat{\Omega})\right)^T \boldsymbol{FIM} \left(\widetilde{\boldsymbol{m}} - \boldsymbol{m}(\hat{\theta}, \hat{\phi}, \hat{\Omega})\right) \tag{2}$$

such that

$$m(\theta, \phi, \Omega) = \left[\langle\mu_x^2\rangle, \langle\mu_y^2\rangle, \langle\mu_z^2\rangle, \langle\mu_x\mu_y\rangle, \langle\mu_x\mu_z\rangle, \langle\mu_y\mu_z\rangle\right]^T, \tag{3}$$

$$\langle\mu_x^2\rangle = \gamma\mu_x^2 + \frac{1-\gamma}{3}, \quad \langle\mu_x\mu_y\rangle = \gamma\mu_x\mu_y, \tag{4}$$

$$\langle\mu_y^2\rangle = \gamma\mu_y^2 + \frac{1-\gamma}{3}, \quad \langle\mu_x\mu_z\rangle = \gamma\mu_x\mu_z, \tag{5}$$

$$\langle\mu_z^2\rangle = \gamma\mu_z^2 + \frac{1-\gamma}{3}, \quad \langle\mu_y\mu_z\rangle = \gamma\mu_y\mu_z, \tag{6}$$

$$\left[\mu_x, \mu_y, \mu_z\right] = [\sin\theta\cos\phi, \sin\theta\sin\phi, \cos\theta], \text{and} \tag{7}$$

$$\gamma = 1 - \frac{3\Omega}{4\pi} + \frac{\Omega^2}{8\pi^2}, \tag{8}$$

where $\gamma$ is the rotational constraint[16] and $\boldsymbol{FIM}$ is the Fisher information (FI) matrix calculated from the basis images. Note that $\widetilde{\boldsymbol{m}}$ and $\boldsymbol{m}$ denote second moment outputs of the maximum likelihood estimator and the weighted least-square estimator respectively. Here, we define the FI matrix associated with estimating the six orientational second moments $\boldsymbol{m}$ as

$$\boldsymbol{FIM} = \sum_{i=1}^{n} \frac{1}{I_i} \nabla I_i^T \nabla I_i, \tag{9}$$

where $i$ denotes the $i^{\text{th}}$ pixel of an image $I \in \mathbb{R}^n$ captured by a camera and $\nabla I_i = \left[\frac{\partial I_i}{\partial m_{xx}}, \frac{\partial I_i}{\partial m_{yy}}, \frac{\partial I_i}{\partial m_{zz}}, \frac{\partial I_i}{\partial m_{xy}}, \frac{\partial I_i}{\partial m_{xz}}, \frac{\partial I_i}{\partial m_{yz}}\right]$. Due to the linearity of the forward imaging model (Eq. 1) in terms of the second moments, the FI matrix can be further simplified as

$$\boldsymbol{FIM} = \sum_{i=1}^{n} \frac{s^2}{I_i} \boldsymbol{B}_i^T \boldsymbol{B}_i. \tag{10}$$

where $\boldsymbol{B}_i$ represents the $i^{\text{th}}$ row of $\boldsymbol{B} \in \mathbb{R}^{n\times 6}$. The weighted least square estimation can be efficiently performed by pre-calculating Hadamard products of each pair of the basis images. The FI matrix assigns weights to each orientational component $m_k$ inversely proportional to the expected measurement variance of the PSFs used in SMOLM. We performed the optimization in Eq. (2) using the $fmincon$ function in MATLAB (Mathworks, R2019a). The eigenvector corresponding to the largest eigenvalue of the second moment matrix[11] was assigned as the initial point of this minimization.

### 6.2 PSF detectability

SMOLM can use any PSF that encodes SM orientation and wobble into its images. The Tri-spot PSF provides highly accurate and precise measurements of SM orientation and wobbling[16] by redistributing the photons from a SM into three spots (in both x- and y-polarized detection channels). It therefore exhibits smaller signal-to-background ratios (SBRs) compared to the standard PSF. In applications with weakly emitting fluorescent molecules or high fluorescence background, a significant fraction or even a majority of SM flashes may not be detectable using the Tri-spot PSF.

Using the Tri-spot PSF, we observed that the average photons detected from single Nile red molecules decreased from 1365±549 (median±std) in a DOPC/DPPC/chol mixture to 883±301 in DOPC/SPM/chol. This decrease could be due to a smaller quantum yield (QY) and a blue-shift in the fluorescence of Nile red in the presence of SPM+chol (QY, 45%; em, 586 nm) compared to DPPC+chol (QY, 60%; em, 595 nm).[17] This observation drove us to choose a different orientation-sensitive PSF with improved SBR for SMOLM imaging in DOPC/SPM/chol lipid membranes.

#### 6.3 Design of the Duo-spot PSF

We therefore designed an orientation-sensitive Duo-spot PSF for the SMOLM imaging of Nile red in DOPC/SPM/chol. The Duo-spot PSF was designed to distinguish the $\boldsymbol{B}_{zz}$ basis image from $\boldsymbol{B}_{xx}$ in the $x$-polarized channel (or $\boldsymbol{B}_{yy}$ in the $y$-polarized channel) for improved polar angle ($\theta$) estimation in SMOLM. In order to achieve this goal, we focused on the intensity distributions of the $\boldsymbol{B}_{yy}$ and $\boldsymbol{B}_{zz}$ images at the back focal plane (BFP) in the $y$ channel (Fig. S7a, the $\boldsymbol{B}_{xx}$ and $\boldsymbol{B}_{zz}$ basis images in $x$-polarized channel are similar but rotated by 90°). The Duo-spot phase mask splits the BFP into two regions as shown in Fig. S7b. Although the Duo-spot PSF exhibits two bright spots in response to an isotropic emitter or a dipole emitter with $\Omega = 2\pi$ sr (Fig. S7c), if the wobbling area $\Omega$ is relatively small (e.g. $\Omega = 0.25\pi$ sr), the brightness ratio between the two spots in each channel changes sensitively in response to the polar angle $\theta$ (Fig. S7d-g). This response essentially produces well-focused, approximately single-spot images when molecules exhibit large or small polar angles ($\theta \approx 0°$ or $\theta \approx 90°$) and allows such molecules to be detected efficiently under low SBR conditions compared to the Tri-spot PSF (Fig. S7f-i). Using our estimation algorithm[14,15] (Supplementary Note 6.1), the Duo-spot PSF yields good estimation precision ($\sigma_\theta^{\text{mean}} = 8.4°$, $\sigma_\phi^{\text{mean}} = 21.8°$, $\sigma_\Omega^{\text{mean}} = 0.16\pi$ sr) and bias ( $|\theta - \theta_0|^{\text{mean}} = 3.5°$, $|\phi - \phi_0|^{\text{mean}} = 0.0°$, $|\Omega - \Omega_0|^{\text{mean}} = -0.11\pi$ sr) for measuring the orientation spectra of fluorescent molecules with a brightness of 950 photons and a background of 5 photons/pixel (Figs. S8 and S9). We utilized the Duo-spot PSF in the SMOLM imaging of SMase-induced lipid composition alteration and domain reorganization (Fig. 3).

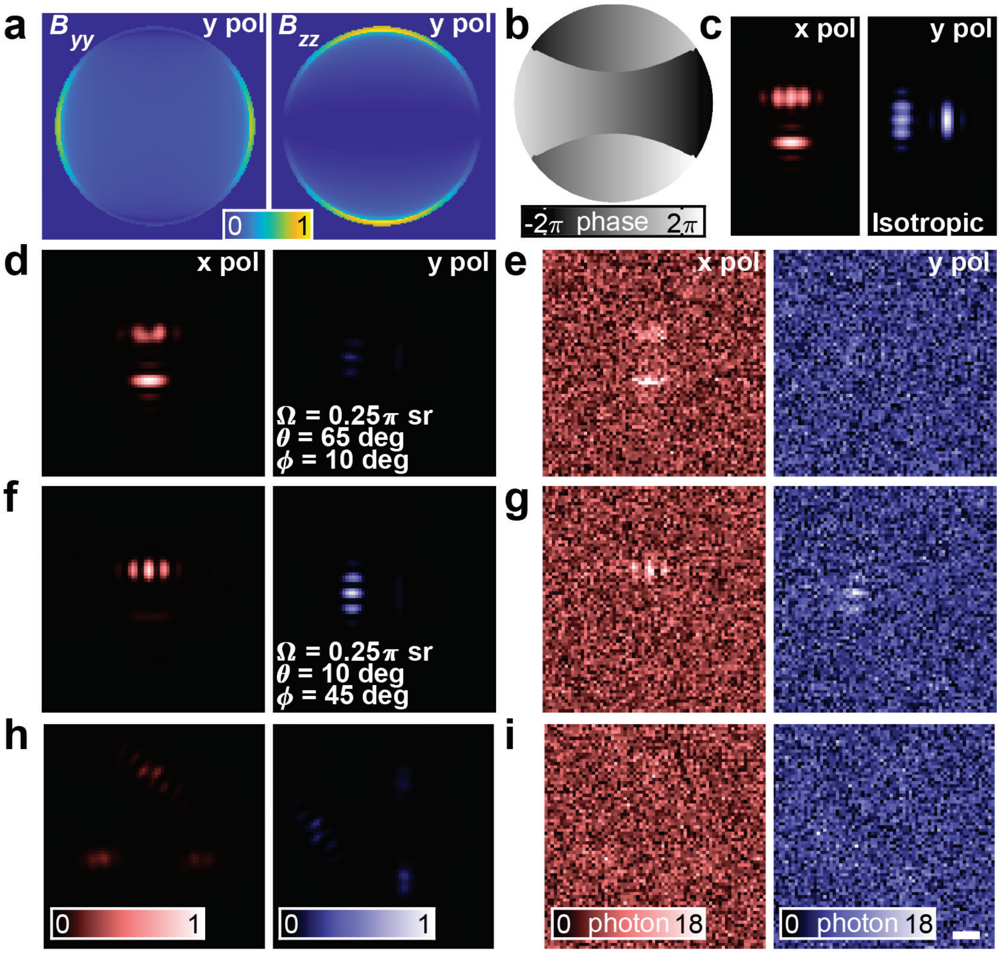


Fig. S7 | Design of the Duo-spot PSF. (**a**) Basis images at the back focal plane corresponding to orientational second-moment components $\langle\mu_y^2\rangle$ and $\langle\mu_z^2\rangle$ in the y-polarized emission channel. Color bar: normalized intensity. (**b**) Duo-spot phase mask. Color bar: phase (rad). (**c**) Simulated Duo-spot PSF for an isotropic emitter. (**d**-**g**) Simulated Duo-spot PSF for a molecule (**d,e**) oriented at $\theta = 65°$, $\phi = 10°$ and wobbling within $\Omega = 0.25\pi$ sr, and a molecule (**f,g**) oriented at $\theta = 10°$, $\phi = 45°$ and wobbling within $\Omega = 0.25\pi$ sr (**d,f**) without and (**e,g**) with Poisson shot noise and background (brightness of 950 photons and background of 5 photons/pixel). (**h,i**) Simulated Tri-spot PSF to a molecule with the same orientation as **f,g**. Color bars: normalized intensity in **d,f,h** and brightness (photon) in **e,g,i**. Scale bar: 500 nm in **c**-**i**.

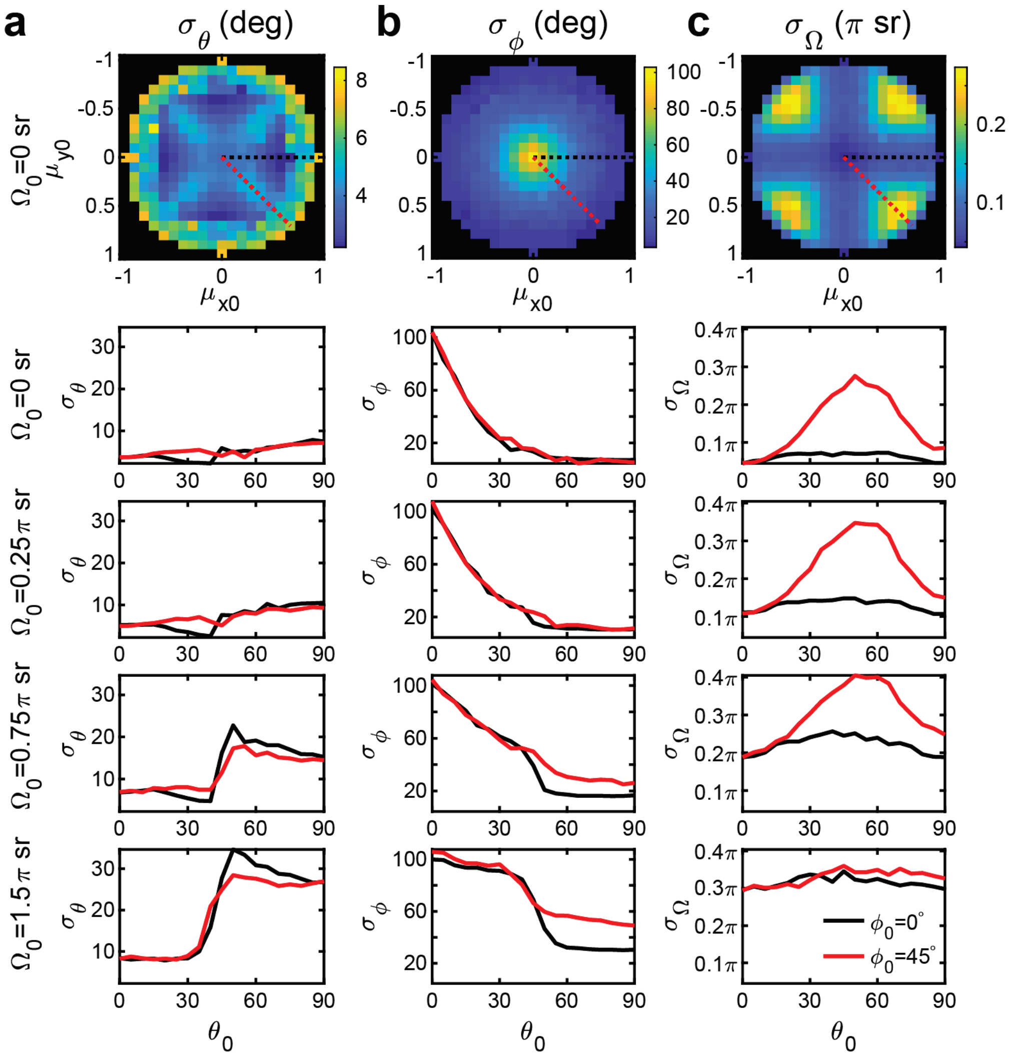


Fig. S8 | Orientation spectra estimation precisions for (**a**) polar angle $\theta$, (**b**) azimuthal angle $\phi$, and (**c**) solid angle $\Omega$ determined from simulated Duo-spot PSF images of dipoles at various orientations. ($\theta_0 = 0° - 90°, \phi_0 = 0° - 360°, \Omega_0 = 0 - 1.5\pi$ sr, $\mu_{x0} = \sin\theta_0 \cos\phi_0, \mu_{y0} = \sin\theta_0 \sin\phi_0$). At each orientation, 1000 independent images were generated with a brightness of 950 photons and a background of 5 photons/pixel. Orientations of simulated molecules were estimated using our maximum-likelihood estimation algorithm, and the orientation estimation precision was computed by taking standard deviation of all estimates at each orientation. Due to symmetry with respect to $\phi_0$ as shown in the first row, the estimation precision is only reported at $\phi_0 = 0°$ (black) and $\phi_0 = 45°$ (red) for $\Omega_0 > 0$ sr.

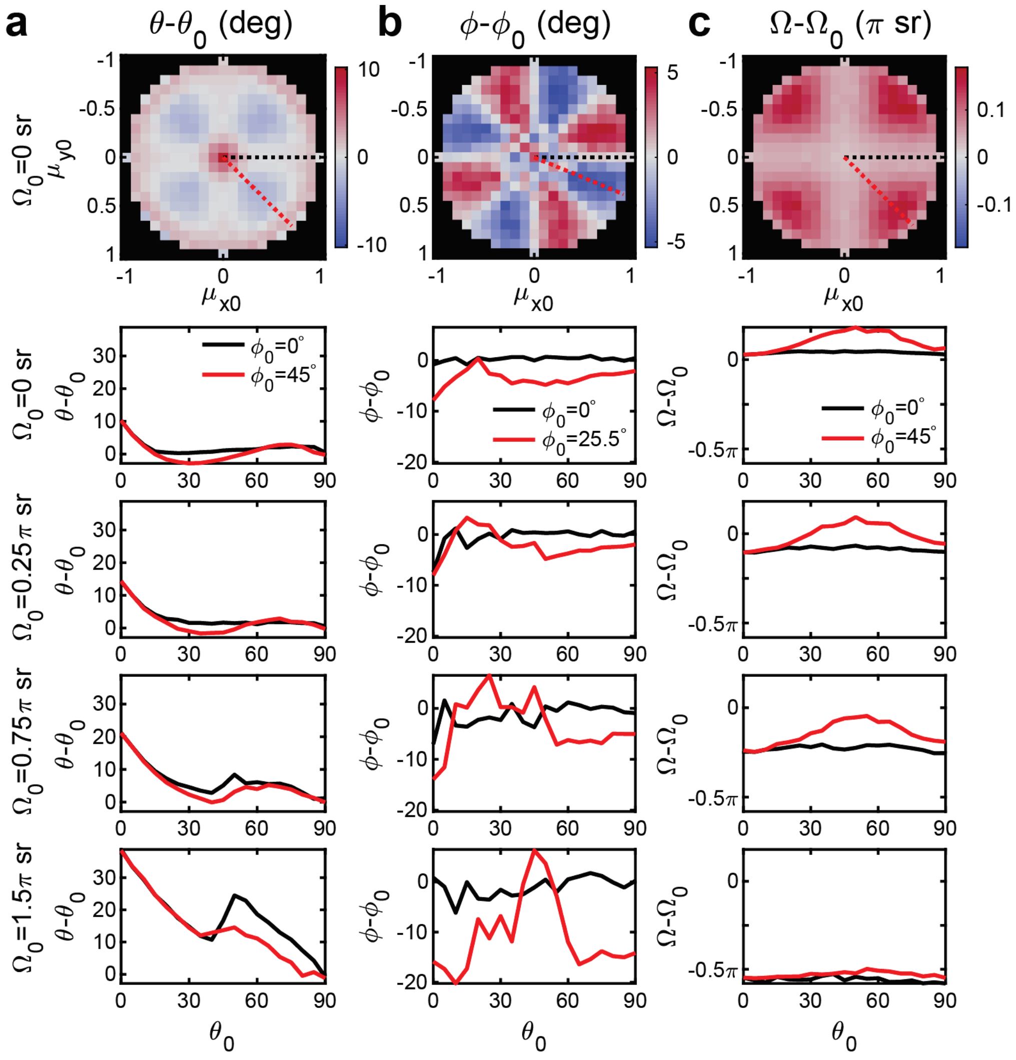


Fig. S9 | Orientation spectra estimation bias for (**a**) polar angle $\theta$, (**b**) azimuthal angle $\phi$, and (**c**) solid angle $\Omega$ determined from simulated Duo-spot PSF images of dipoles at various orientations . ($\theta_0 = 0° - 90°, \phi_0 = 0° - 360°, \Omega_0 = 0 - 1.5\pi$ sr, $\mu_{x0} = \sin\theta_0 \cos\phi_0$, $\mu_{y0} = \sin\theta_0 \sin\phi_0$ ). At each orientation, 1000 independent images were generated with a brightness of 950 photons and a background of 5 photons/pixel. Orientations of simulated molecules were estimated using our maximum-likelihood estimation algorithm, and the orientation estimation bias was computed by averaging all measurement deviations at each orientation. Due to symmetry with respect to $\phi_0$ as shown in the first row, the estimation bias is only reported at $\phi_0 = 0°$ (black), 45° (red, for polar and solid angles) or 22.5° (red, for azimuthal angle) for $\Omega_0 > 0$ sr.

#### 6.4 SMOLM performance using the Duo-spot PSF for measuring chol in SLBs

Experimentally, we used the Duo-spot PSF to image the orientation spectra of Nile red in the presence of increased chol. As shown in Fig. S10a, both the polar angle and solid angle of Nile red decrease when the chol concentration increases from 0% to 40% in SPM. This trend matches our observations of Nile red in SPM using the Tri-spot PSF (Fig. S10b).

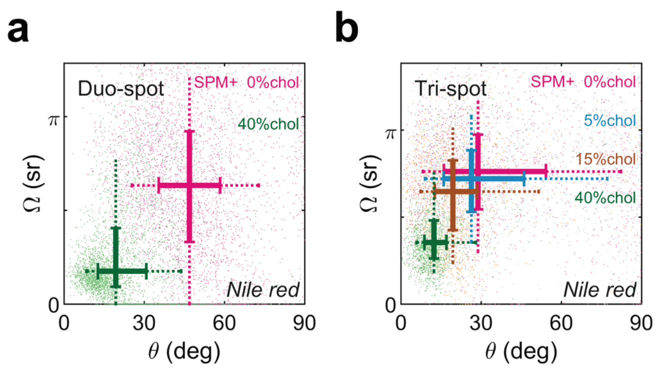


Fig. S10 | Orientation (polar angle θ) and wobble (solid angle Ω) of Nile red in SPM with various cholesterol levels from 0% to 40% measured using the (**a**) Duo-Spot PSF and (**b**) Tri-spot PSF.

Fifty representative frames of raw SMOLM images corresponding to the data in Fig. S10 (SPM and SPM+40% chol SLBs using Tri-spot PSF and Duo-spot PSF) are shown in Movie S2.

### Supplementary Note 7. Molecular lateral diffusion distorts single molecule PSFs and biases orientation and wobble estimation

In the liquid phase, lipid acyl chains are kinked and loosely packed, causing fluorescence probes embedded within the lipid bilayer to diffuse rapidly (D=2~5 μm$^2$/s)[18]. This lateral diffusion results in an average $4\times10^2$-$7\times10^2$ nm ($\sqrt{4Dt}$) displacement within a 30-ms camera exposure time. This distance is of the same magnitude as the size of an optical PSF. Therefore lateral diffusion could distort the Tri-spot or Duo-spot PSF and potentially bias SMOLM orientation estimates.

To quantitatively understand how the diffusion-induced PSF distortion influences orientation estimation using different orientation-sensitive PSFs, we used a random walk to simulate the lateral trajectory of single molecule within one camera frame and thus generate the PSF image. A brightness of 1365 photons and background of 11 photons/pixel were used to match our typical lipid imaging conditions. The lateral diffusion coefficients were set to 5 μm$^2$/s and 0.005 μm$^2$/s for liquid and gel phases, respectively. Each trajectory within one camera frame (30 ms) contains 100 steps of random walk, and for each set of ground truth polar angle θ, azimuth angle ϕ, and solid angle Ω, we simulated 100 independent camera frames. Finally, the orientation and wobble of simulated PSFs were estimated using the same maximum-likelihood estimation algorithm (Supplementary Note 6.1) as used for analyzing experimental data.

In the gel phase, the average lateral displacement of a single probe molecule within 30-ms camera exposure time is 25 nm ($\sqrt{4Dt}$). Our simulation results indicate that both the Tri-spot and Duo-spot PSFs show accurate orientation (polar angle and solid angle) estimates (blue and light blue plots in Fig. S11). In the liquid phase, however, the lateral diffusion-induced PSF distortion causes large biases of both PSFs (red and orange plots in Fig. S11). Especially for ground truth orientations with small polar angle ($\theta<5°$) and wobble ($\Omega<0.25\pi$ sr), the polar angle was overestimated by up to 20 degrees using the Duo-spot PSF compared to the Tri-spot PSF.

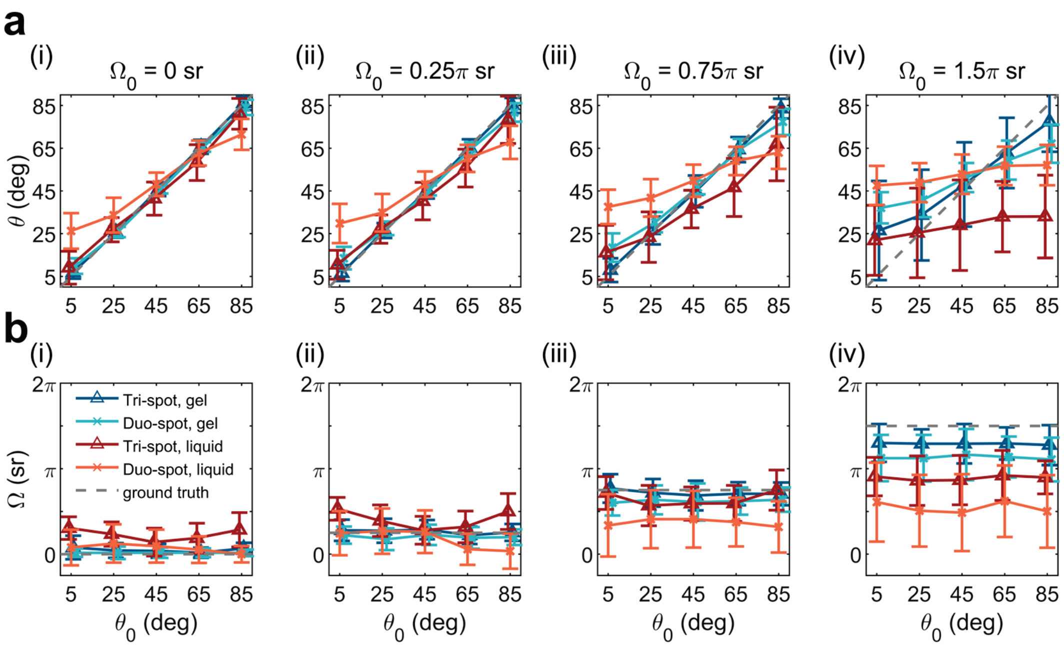


Fig. S11 | (**a**) The estimated orientation (polar angle, θ) and (**b**) wobble (solid angle Ω) for molecules with various degrees of wobbling: ground truth solid angles $\Omega_0$ of (**i**) 0, (**ii**) 0.25π, (**iii**) 0.75π, (**iv**) 1.5π sr in gel (blue or light blue) and liquid (red or orange) phase of lipid membrane using the Tri-spot (△) and Duo-spot (×) PSFs. The gray dashed line indicates the ground truth. The diffusion coefficients are 0.005 $\mu m^2/s$ in gel phase and 5 $\mu m^2/s$ in liquid phase. $\Omega_0$: ground truth solid angle; $\theta_0$: ground truth polar angle.

Although the Duo-spot was rationally designed for measuring out-of-plane orientations precisely and its performance on orientation detection has been detailed characterized in the previous section (Supplementary Note 6), our simulation results indicate its performance is more easily affected by the lateral diffusion-induced PSF distortion in liquid phase of lipid bilayer than Tri-spot. Currently, we did not attempt to compensate for estimation bias caused by lateral diffusion within our maximum-likelihood estimation algorithm. Any orientation-sensitive PSFs used for SMOLM could suffer varying degrees of estimation bias due to fluorophore diffusion.

Practically, in SMOLM lipid imaging, as long as the apparent orientation and wobble angles show separable changes and these changes can be interpreted using the measurements obtained from single-phase lipid samples, both the Duo-spot or Tri-spot PSF remain powerful tools for separating lipid domains and detecting the compositional alternations in lipid membrane.

### Supplementary Note 8. Comparison of the Duo-spot vs. Tri-spot PSFs for SMOLM imaging of DOPC/SPM/chol SLBs

For SMOLM imaging of mixed DOPC/SPM/chol SLBs (Fig. 3), we used the Duo-spot PSF due to its improved SBR over the Tri-spot PSF. Although the SMOLM images (polar angle map, solid angle map, and phase-index map) resolve the Lo/Ld domains well (Fig. 3), the measured or apparent orientation spectra using the Duo-spot PSF are different from those of the Tri-spot PSF. For identical SLB compositions of DOPC/SPM/chol (35:35:30, molar ratio), the measured polar angle of Nile red shows a significant difference when using the Tri-spot PSF (17.6±16.8°, median±std) vs. the Duo-spot PSF (40.0±12.9°), as shown in Fig. S12a. The measured wobble (solid angle) shows less variation between PSFs (Fig. S12b): 0.62π±0.31π sr (Tri-spot PSF) vs. 0.60π±0.41π sr (Duo-spot PSF).

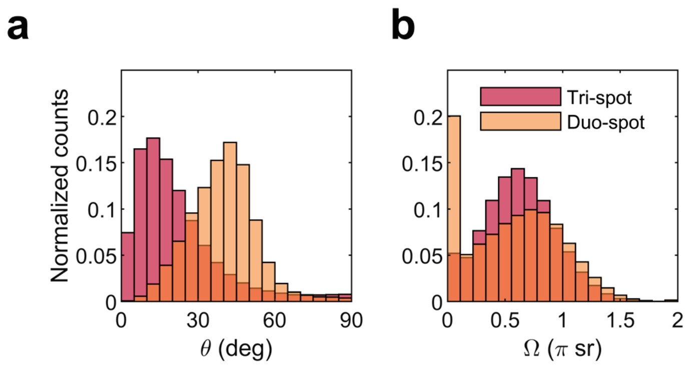


Fig. S12 | (**a**) Orientation (polar angle θ) and (**b**) wobble (solid angle Ω) of Nile red within a mixture of DOPC/SPM/chol (35:35:30, molar ratio), measured using the Tri-spot PSF (red) and Duo-spot PSF (orange).

The lipid mixture DOPC/SPM/chol contains both Lo and Ld phases with different lipid compositions and different diffusion coefficients. To better understand the observed discrepancies in Fig. S12, it is reasonable to compare the orientation spectra of Nile red in single-phase lipid samples, e.g. DOPC(+chol) SLB, SPM+chol SLB, and SPM+cer SLB.

In the single-phase lipid samples, minor differences were observed within the Nile red orientation spectra in SPM+chol and SPM+cer phases when comparing the Duo-spot (Fig. S13b, green and purple plots) to the Tri-spot PSF (Fig. S13a, green and purple plots). These phases have low diffusion coefficients (0.10±0.02 μm$^2$/s for Lo phase[19], 0.005 μm$^2$/s for gel phase), and the molecules show small lateral displacement (< 100 nm) within our typical camera exposure time. The estimated orientation and wobble using Tri-spot and Duo-spot approximately match each other, as shown in our simulations of diffusing molecules (Fig. S11).

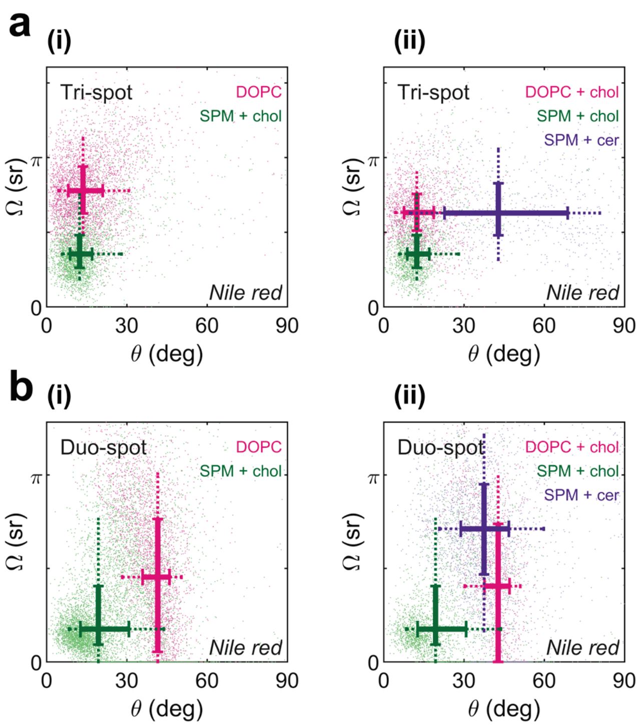


Fig. S13 | Orientation (polar angle θ) and wobble (solid angle Ω) of Nile red in the single-phase SLBs consisting of (**i**) DOPC and SPM+chol, and (**ii**) DOPC+chol, SPM+chol, and SPM+cer, imaged using the (**a**) Tri-spot PSF and (**b**) Duo-spot PSF.

However, within the DOPC Ld phase, major differences were observed in the estimated polar angle between the Tri-spot PSF (13.6±11.6°, Fig. S13a(i), pink plot) and the Duo-spot PSF (41.5±8.5°, Fig.

S13b(i), pink plot). Similar differences were also observed in the DOPC+chol Ld phase (Fig. S13a(ii) and Fig. S13b(ii), pink plots). We therefore investigated how molecular diffusion affects the estimated orientation and wobble in DOPC(+chol) SLBs. For our simulations of diffusive molecules, we used the typical brightness, background, diffusion coefficient (5 $\mu m^2/s$), and exposure time that match experimental conditions.

As shown in Table S1, for the molecule with a ground-truth orientation of 16° (polar angle) and wobble of π sr (solid angle) in DOPC Ld SLBs, the estimated polar angle when using the Tri-spot PSF from simulated images (13.6±6.0°) matches the experimental result (13.6±11.6°) very well. However, the Duo-spot simulation reports the polar angle as 33.2±8.23°, which is about 20° larger than that measured by the Tri-spot experimentally. Although our Duo-spot simulation does not perfectly match the measured experimental polar angle (41.5±8.5°), our simulations confirm that polar angles are more likely to be overestimated by using the Duo-spot PSF, compared to Tri-spot; this overestimate is likely caused by high lateral diffusion and PSF distortion in Ld phases. On the other hand, the estimated solid angle in simulated images (0.95π±0.05π sr) is much larger than experimental observations (0.46π±0.40π sr) for the Duo-spot PSF and is likely caused by some other form of imaging model-experimental mismatch. Similar simulation results were obtained for the DOPC+chol sample (Table S1); the Tri-spot PSF experimental measurements largely match those from simulated images of diffusive molecules, while the experimental orientation measurements using the Duo-spot PSF are systematically larger than those from simulated images.Although our simulations of laterally diffusing molecules partially explain discrepancies in the orientation spectra in single-phase lipid samples between the Duo-spot and Tri-spot PSFs, the orientation spectra are still well separable in DOPC(+chol), SPM+chol, and SPM+cer SLBs (Fig. S13). Therefore, both the Tri-spot and Duo-spot PSFs are able to distinguish reliably various lipid phases in lipid mixtures of DOPC/SPM/chol.

Table S1. Comparison of simulated and experimental SMOLM orientation measurements (polar angle θ and wobble solid angle Ω) (median±std) of laterally diffusing Nile red in single-phase lipid samples using the Tri-spot and Duo-spot PSFs.

| Single-phase lipid sample | | DOPC SLB | | DOPC+chol SLB | |
|---|---|---|---|---|---|
| | | Polar angle, θ (deg) | Solid angle, Ω (π sr) | Polar angle, θ (deg) | Solid angle, Ω (π sr) |
| Ground truth for simulation | | 16 | 1 | 12 | 0.74 |
| Tri-spot PSF | experiment | 13.6±11.6 | 0.78±0.24 | 12.3±9.1 | 0.63±0.20 |
| | simulation | 13.6±6.0 | 0.76±0.01 | 13.0±8.9 | 0.64±0.01 |
| Duo-spot PSF | experiment | 41.5±8.5 | 0.46±0.40 | 42.8±8.0 | 0.41±0.39 |
| | simulation | 33.2±8.3 | 0.95±0.05 | 26.3±8.1 | 0.84±0.06 |

### Supplementary Note 9. SMLM of SMase-induced lipid domain reorganization

With high doses of SMase (500 mU/mL) applied to DOPC/SPM/chol (35:35:30, molar ratio) SLBs, we observed extensive changes in the morphologies of Lo domains (dark regions in Fig. S14) due to the enzymatic generation of ceramide. The total Lo domain size reduces from 32.2 μm$^2$ (Fig. S14a) to 15.9 μm$^2$ (Fig. S14b). On average, the total Lo domain area decreases by 50%, which is mainly caused by tightly packed ceramide that decreases the distance between the lipid molecules via extensive hydrogen bonding.[20,21]

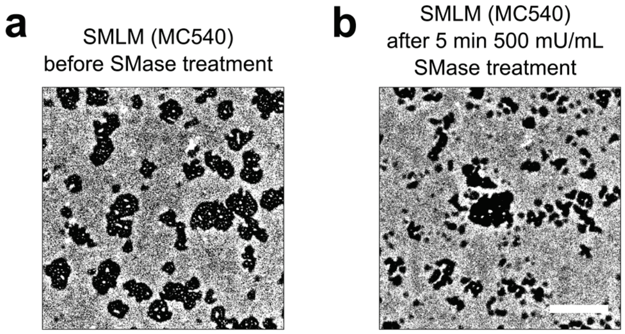


Fig. S14 | MC540 SMLM images show Lo (dark) and Ld (bright) domains (**a**) before and (**b**) after 5 min of 500 mU/mL SMase treatment in DOPC/SPM/chol (35:35:30, molar ratio) SLBs. Scale bar: 2 μm. Bin size: 28 nm.

### Supplementary Note 10. Principal component analysis (PCA) analysis of SMOLM data

#### 10.1 PCA analysis to distinguish gel (or Lo) and liquid (or Ld) phases

For a lipid mixture containing gel (or Lo) and liquid (or Ld) phases, we performed principle component analysis (PCA) on SMOLM datasets containing orientation (polar angle θ) and wobble (solid angle Ω) measurements. The data of θ and Ω were first standardized by removing the mean and scaling to unit variance, and subsequently, PCA was applied to reduce the dimensionality.[22] We designated the resulting PCA scores (first component) as the phase index for generating SMOLM phase-index maps (Fig. 2, S4, and S5).

Gel (or Lo) and liquid (or Ld) phases can be discriminated by using the localization density (per pixel) in SMLM. For example, in the lipid mixture of DOPC/DPPC/chol in Fig. 2b, regions with fewer than 4 localizations/bin in the SMLM image are assigned as the Lo phase, and regions with counts above 4 per pixel are designated as the Ld phase. Using these classifications as a ground-truth calibration for our phase-index values (arb. units), we compute the distribution of phase index in each phase (Lo: -0.63±0.91, Ld: 0.06±1.02), and set the midpoint between the mean indices (-0.28, arb. units) as the threshold to separate Lo and Ld domains.

#### 10.2 KPCA analysis to distinguish lipid phases in DOPC/SPM/chol with SMase treatment

##### 10.2.1 Analysis procedures

In the lipid mixture of DOPC/SPM/chol after SMase treatment, three different phases could coexist: a DOPC (or DOPC+chol) Ld phase, an SPM+chol Lo phase, and an SPM+cer Lo phase. The orientation spectra of Nile red are able to distinguish these three phases in single-phase samples (Fig. S13b). In order to generate phase-index maps in mixed samples, we applied support vector machines (SVM) and kernel principal component analysis (KPCA) to conduct non-linear dimensionality reduction on the SMOLM data (polar angle θ and solid angle Ω) via the following procedures:[22]

1). Use SMOLM data of single-phase lipid samples (DOPC SLB, SPM+chol SLB, and SPM+cer SLB) as training data to fit the SVM model.

The performance of SVM model fitting was tested by performing classification on the same training data. We found that using the six orientational second moments[16] instead of polar and solid angles from SMOLM produced higher accuracy scores and better classification results (Fig. S15). We therefore used the second moment estimates from our maximum likelihood estimator (Supplementary Note 6.1) for all SVM and KPCA analyses.

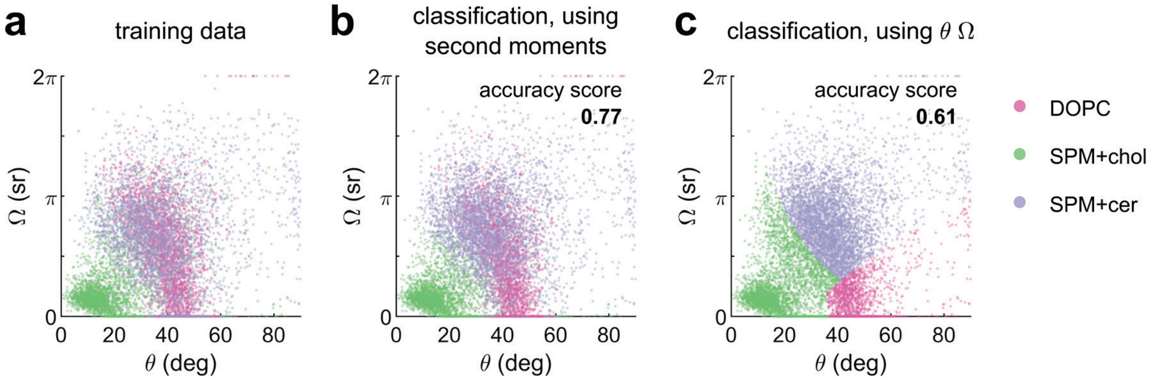


Fig. S15 | (**a**) Orientation (polar angle θ) and wobble (solid angle Ω) of Nile red in the single-phase lipid samples consisting of DOPC, SPM+chol, and SPM+cer using the Duo-spot PSF. Each sample contains 3300 data points, and total 9900 data points were used for SVM training. (**b**,**c**) SVM classification results on the same training data using (**b**) six second moments and (**c**) polar and solid angles.

2). Use the SVM model to classify the SMOLM results of DOPC/SPM/chol after SMase treatment ($t_3$ in Fig. 3) into three classes (DOPC, SPM+chol, SPM+cer).

The purpose of this step is to classify and label the experimental data, and select balanced data from three phases to fit the KPCA model in the next step.

3). Select equal numbers (11833 data points for each class) of data points from the three classes in step 2, and fit to a KPCA model (using rbf kernel).

4). Apply this KPCA model in step 3 to transform entire SMOLM dataset ($t_0$,$t_1$,$t_2$,$t_3$ in Fig. 3), and use the KPCA score of the first component as the SMOLM phase index.

The SMOLM phase-index maps (Fig. 3, S16a(iv), S17b) during SMase treatment were all generated using above procedures.

#### 10.2.2 Analyzing single-phase lipid samples

We calculated the phase-index of the training data (single-phase lipid samples, Fig. S13b), including SPM+chol, SPM+cer, DOPC, and DOPC+chol, using the procedures above. The results are summarized in Table S2.

Nile red in SPM+chol (Lo phase) exhibits small polar and solid angles with negative phase indices (Table S2). Both the orientation spectra (polar and wobble angles) and phase index increase as the lipid composition changes to a SPM+cer Lo phase (Table S2). Therefore, Nile red SMOLM imaging has excellent sensitivity to distinguish the conversion from chol-rich to ceramide-rich Lo domains. In Ld phases (DOPC or DOPC+chol), the ordering effect of chol is weak. The addition of chol does not have a significant impact on the observed Nile red orientation spectra and phase indices (Table S2), which is similar to observations using the Tri-spot PSF in Fig. S2b. Therefore, we do not discriminate between DOPC and DOPC+chol domains when imaging enzyme-mediated changes to lipid composition (Fig. 3).

Table S2. SMOLM measurements of the orientation (polar angle, θ), wobble (solid angle, Ω), and phase index (median±std) of Nile red in single-phase lipid samples using the Duo-spot PSF.

| Single-phase lipid sample | SPM+chol SLB | SPM+cer SLB | DOPC SLB | DOPC+chol SLB |
|---|---|---|---|---|
| Polar angle, θ (deg) | 19.4±14.1 | 37.5±15.7 | 41.5±8.5 | 42.8±8.0 |
| Solid angle, Ω (π sr) | 0.18±0.29 | 0.71±0.37 | 0.46±0.40 | 0.41±0.39 |
| Phase-index (KPCA score, arb. units) | -0.070±0.302 | 0.042±0.271 | 0.069±0.258 | 0.080±0.284 |

We further calculated the mean value of phase index in single-phase lipid samples of SPM+chol SLBs and SPM+cer SLBs. The midpoint between these means (-0.014, arb. units) was used to determine the phase-index threshold to separate the SPM+chol and SPM+cer phases in Fig. 3c(iv), Fig. 3e(iii), and Fig. S17b.

### Supplementary Note 11. SMOLM data visualization using the Duo-spot PSF

When using the Tri-spot PSF, we use the median value of the localizations in each bin to generate the SMOLM maps (polar angle map, solid angle map and phase-index map), as stated in the Methods section. However, when using this same criterion for orientation spectra captured by the Duo-spot PSF, we noticed that many "small-sized" Lo regions with small solid angles appear in SMOLM solid angle maps (Fig. S16a(ii), Fig. S16b(ii)). These discrepancies in the SMOLM maps are caused by variations in the orientation spectra measured by the Duo-spot and Tri-spot PSFs (Supplementary Note 8, Fig. S12).

To analyze these regions in more detail, we applied $\Omega$ and object size thresholds to create three regions (magnified view in Fig S16b). Region 1 contains all bins with $\Omega > 0.4\pi$ sr (median value in Lo phase). Region 3 contains large objects (object size > 8 contiguously connected bins) with $\Omega \leq 0.4\pi$ sr. We assign the remaining bins to Region 2 that contains relatively sparse and small objects (object size $\leq$ 8 bins) with small $\Omega$ ($\leq 0.4\pi$ sr). We analyze the polar angle, solid angle, and phase index from all localizations within each region (Fig. S16b). The distribution of phase index (Fig. S16c(iii)) and polar angle (Fig. S16c(ii)) show that regions 1 and 2 contain lipids in the Ld phase; their distributions of polar angle and phase index all overlap one another. Conversely, these data suggest that region 3 belongs to the Lo phase.

However, the solid angle histogram (Fig. S16c(i)) indicates that region 2 contains populations of both small $\Omega$ (as in region 3) and large $\Omega$ (as in region 1). As shown in Fig. S16d(i), for percentiles below 55, region 2 appears to be same phase as region 3 (Lo phase), which is inconsistent with the polar angle and phase-index data (Fig. S16d(ii)(iii)). In contrast, the solid angles at the $75^{th}$ percentile or above for regions 1 and 2 are in good agreement. Therefore, we chose the $75^{th}$ percentile to represent the SMOLM maps of solid angle and polar angle for all Duo-spot PSF measurements, while keep using the $50^{th}$ percentile for the phase-index map. This choice creates consistent visualizations of Lo and Ld domains in all SMOLM maps (Fig. 3b).

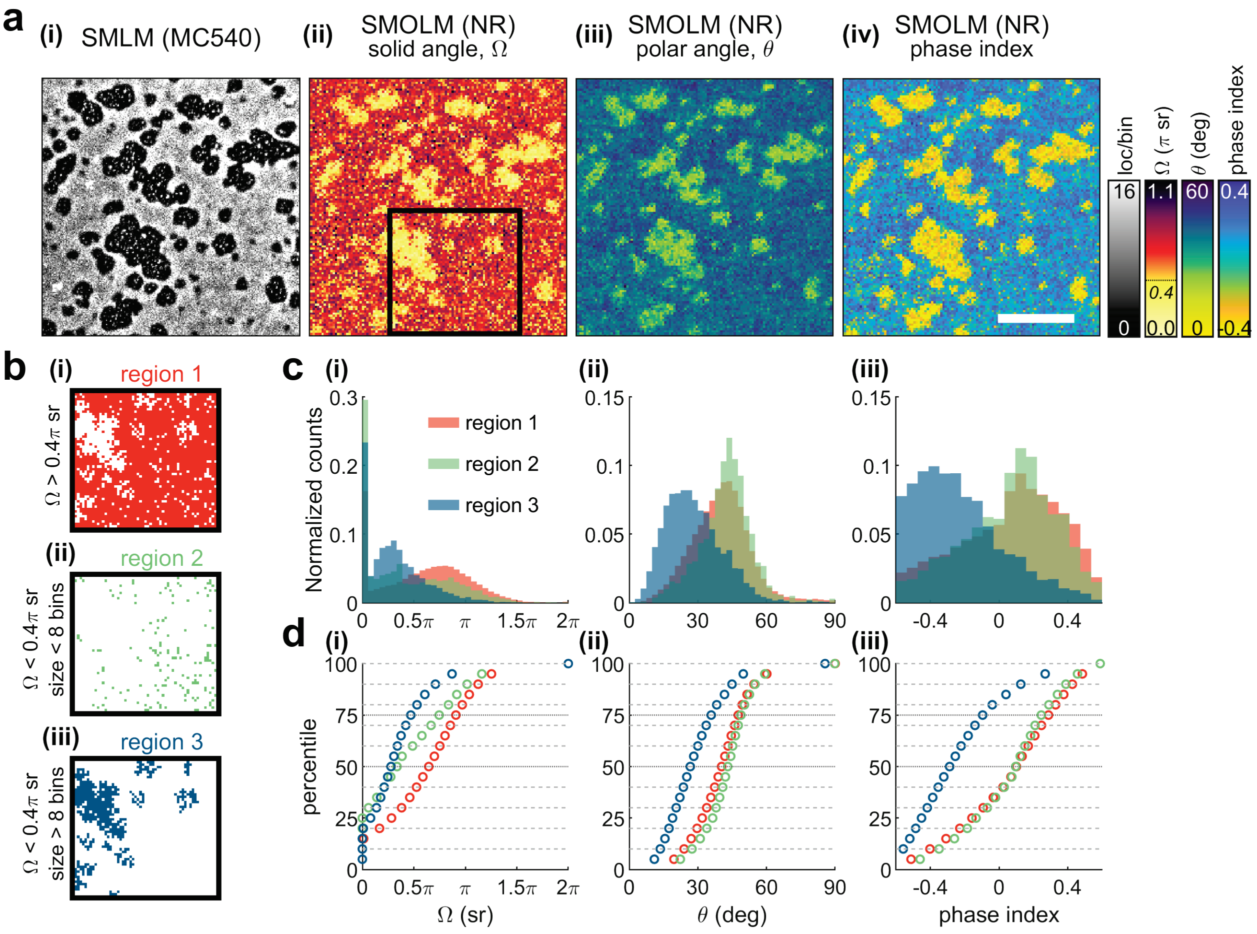


Fig. S16 | (**a**) (**i**) Conventional MC540 SMLM image and (**ii-iv**) SMOLM images ((**ii**) solid angle (Ω), (**iii**) polar angle (θ), (**iv**) phase-index map) of Nile red within a mixed DOPC/SPM/chol (35:35:30, molar ratio) SLB. The SMOLM maps were generated using the median value of localizations within each bin. (**b**) Magnified view of the boxed region (3.22×3.39 μm$^2$) in **a**. (**i-iii**) Three regions selected using Ω and object size thresholds. (**c**) Histograms of (**i**) solid angle, (**ii**) polar angle, and (**iii**) phase index of all localizations within each region. (**d**) Cumulative distribution functions of the measured (**i**) solid angle, (**ii**) polar angle, and (**iii**) phase index from all populations within each region. Scale bar: 2 μm in **a**. Bin size: 28 nm in SMLM, 58.5 nm in SMOLM and **b**.

### Supplementary Note 12. SMase-mediated lipid composition changes in Lo domains

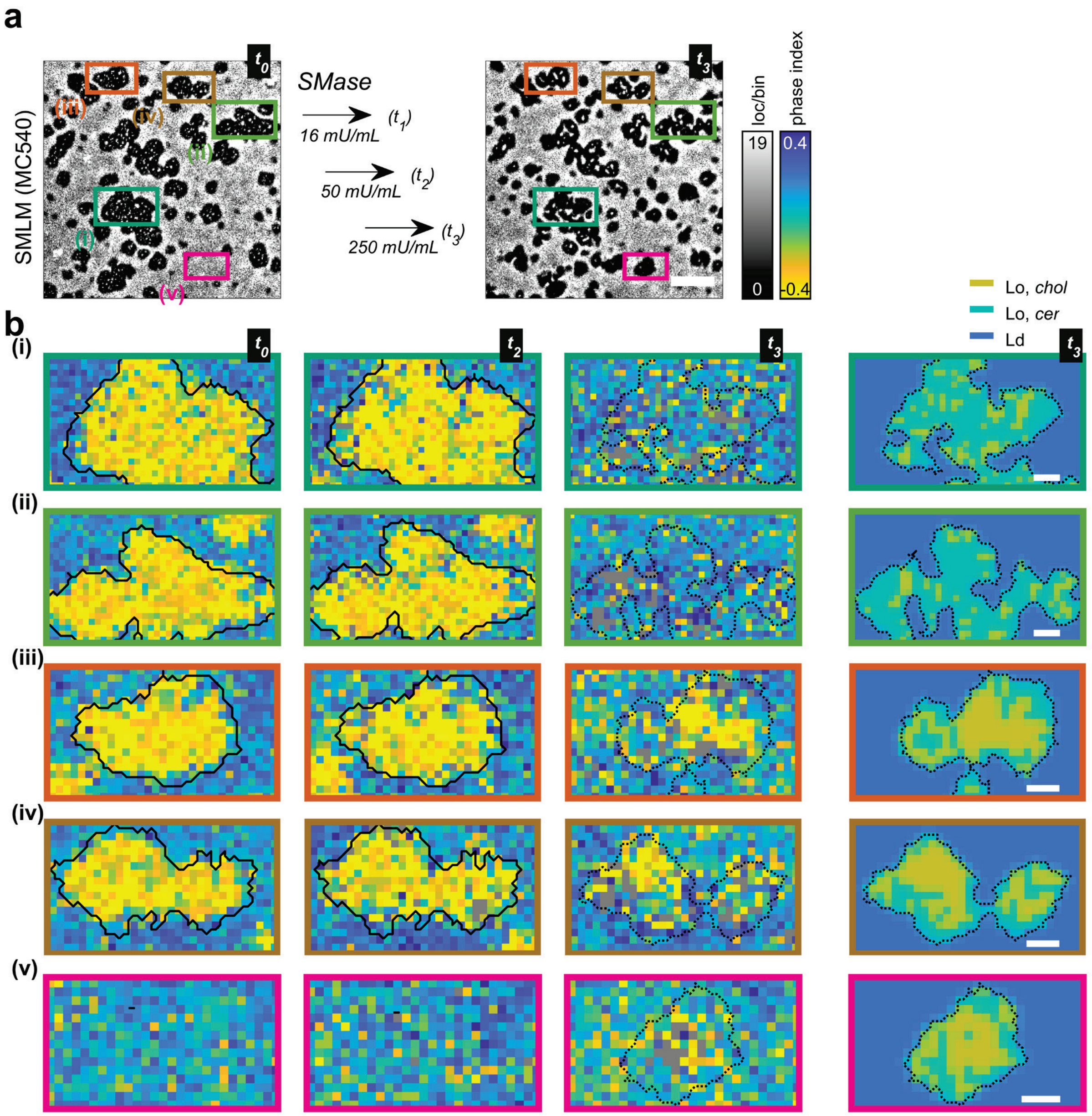


Fig. S17 | (**a**) Conventional MC540 SMLM images of DOPC/SPM/chol before ($t_0$) and after ($t_3$) three doses of SMase treatment. (**b**) Magnified view of SMOLM Nile red phase-index map from (**i-v**) five boxed regions in **a** at $t_0$, $t_2$, $t_3$, and lipid composition map at $t_3$, indicating the lipid composition changes of the existing and newly generated Lo domain by SMase. Scale bar: 2 μm in **a**, 200 nm in **b**. Bin size: 28 nm in SMLM, 45 nm in SMOLM.

The activity of SMase (conversion of SPM to ceramide) varies among different Lo domains. Analysis on multiple single Lo domains (Fig. S17a) reveals both nearly complete conversion (Fig. S17b(i)(ii)) and partial conversion (Fig. S17b(iii)(iv)) to cer-rich SPM phases. There also exist newly formed Lo domains (Fig. S17b(v)) containing well-separated chol-rich and cer-rich phases.

### Supplementary Note 13. Orientation spectra statistics of single-phase SLBs

Table S3. SMOLM orientation spectra statistics and experimental conditions for single-phase lipid samples imaged using the Tri-spot PSF.

| Single-phase lipid sample | Polar angle, θ (deg) | Solid angle, Ω (π sr) | Phase-index (arb. units) | SM brightness (photon) | Background (photon/pixel) | Exposure time (ms) | Figure |
|---|---|---|---|---|---|---|---|
| *DiI* | | | | | | | |
| DPPC | 73.6 ± 21.2 | 0.21 ± 0.41 | -- | 19311.3 | 36.9 | 200 | Fig. 1c |
| *MC540* | | | | | | | |
| DOPC | 17.5 ± 14.2 | 0.71 ± 0.22 | -- | 1369.1 | 4.6 | 30 | Fig. 1d, S5b, S6a |
| DPPC | 73.1 ± 23.7 | 0.68 ± 0.41 | -- | 4334.5 | 15.3 | 100 | Fig. S5b |
| DPPC+40% chol | 15.2 ± 17.8 | 0.60 ± 0.32 | -- | 1963.5 | 15.6 | 100 | Fig. S6a |
| *Nile red (NR)* | | | | | | | |
| DPPC | 26.0 ± 19.2 | 0.96 ± 0.31 | -- | 6033.1 | 13.6 | 100 | Fig. 1e |
| DPPC+5% chol | 15.5 ± 19.0 | 0.82 ± 0.31 | -- | 4046.8 | 14.1 | 100 | Fig. 1e |
| DPPC+10% chol | 12.8 ± 16.8 | 0.63 ± 0.29 | -- | 5018.5 | 14.1 | 100 | Fig. 1e |
| DPPC+20% chol | 11.0 ± 13.2 | 0.32 ± 0.21 | -- | 3151.3 | 13.8 | 100 | Fig. 1e |
| DPPC+40% chol | 8.7 ± 7.6 | 0.26 ± 0.18 | -0.63 ± 0.91 | 4242.0 | 14.0 | 100 | Fig. 1e, 1f, S6b |
| DOPC+40% chol | 15.1 ± 11.8 | 0.73 ± 0.21 | -- | 1614.2 | 4.4 | 30 | Fig. 1f, S2b, S13a |
| POPC+40% chol | 13.8 ± 10.2 | 0.56 ± 0.21 | -- | 1276.0 | 4.7 | 30 | Fig. 1f, S2a |
| DPPC + MβCD-chol 10 μM 10 min | 24.8 ± 17.3 | 0.68 ± 0.33 | -- | 5738.9 | 13.3 | 100 | Fig. S1a |
| + MβCD-chol 40 μM 3 min | 15.2 ± 18.0 | 0.31 ± 0.28 | -- | 5576.3 | 13.3 | 100 | Fig. S1a |
| + MβCD-chol 40 μM 3 min | 9.8 ± 13.4 | 0.23 ± 0.20 | -- | 5311.4 | 14.1 | 100 | Fig. S1a |

| Single-phase lipid sample | Polar angle, θ (deg) | Solid angle, Ω (π sr) | Phase-index (arb. units) | SM brightness (photon) | Background (photon/pixel) | Exposure time (ms) | Figure |
|---|---|---|---|---|---|---|---|
| + MβCD-chol 80 μM 3 min | 8.4 ± 8.2 | 0.20 ± 0.14 | -- | 5251.1 | 13.4 | 100 | Fig. S1a |
| + MβCD-chol 40 μM 3 min | 7.7 ± 5.9 | 0.28 ± 0.09 | -- | 5467.8 | 13.8 | 100 | Fig. S1b |
| + MβCD-chol 80 μM 3 min | 7.5 ± 4.2 | 0.27 ± 0.09 | -- | 5214.4 | 13.6 | 100 | Fig. S1b |
| + MβCD-chol 120 μM 5 min | 6.5 ± 2.8 | 0.26 ± 0.08 | -- | 5255.0 | 13.8 | 100 | Fig. S1b |
| + MβCD-chol 400 μM 10 min | 7.0 ± 3.1 | 0.28 ± 0.07 | -- | 5017.8 | 13.5 | 100 | Fig. S1b |
| DPPC+40% chol+4% mel | 10.8 ± 10.2 | 0.28 ± 0.21 | -- | 2501.9 | 7.6 | 50 | Fig. S1c |
| DPPC+40% chol+30% mel | 17.9 ± 18.4 | 0.39 ± 0.31 | -- | 1775.9 | 6.1 | 30 | Fig. S1c |
| POPC | 17.4 ± 15.1 | 0.72 ± 0.25 | -- | 1377.3 | 4.5 | 30 | Fig. S2a |
| POPC+10% chol | 18.4 ± 13.9 | 0.71 ± 0.24 | -- | 1390.5 | 4.5 | 30 | Fig. S2a |
| POPC+20% chol | 16.1 ± 12.9 | 0.68 ± 0.22 | -- | 1341.8 | 4.4 | 30 | Fig. S2a |
| DOPC | 13.6 ± 11.6 | 0.78 ± 0.24 | 0.06 ± 1.02 | 2015.6 | 4.5 | 30 | Fig. S2b, S6b, S13a |
| DOPC+10% chol | 13.5 ± 12.9 | 0.78 ± 0.21 | -- | 1828.0 | 4.4 | 30 | Fig. S2b |
| DOPC+20% chol | 17.0 ± 15.5 | 0.82 ± 0.22 | -- | 1531.7 | 4.4 | 30 | Fig. S2b |
| SPM | 28.8 ± 25.9 | 0.76 ± 0.32 | -- | 1721.9 | 9.5 | 100 | Fig. S2c |
| SPM+5% chol | 26.3 ± 23.3 | 0.72 ± 0.28 | -- | 1673.7 | 9.2 | 100 | Fig. S2c |
| SPM+15% chol | 19.4 ± 18.3 | 0.65 ± 0.28 | -- | 1824.1 | 9.0 | 100 | Fig. S2c |
| SPM+40% chol | 12.3 ± 11.1 | 0.36 ± 0.23 | -- | 1817.2 | 8.7 | 100 | Fig. S2c, S13a |
| SPM+cer | 42.8 ± 25.5 | 0.63 ± 0.28 | -- | 3654.7 | 11.8 | 100 | Fig. S13a |
| *Nile blue* | | | | | | | |
| DPPC | 33.3 ± 25.5 | 0.88 ± 0.33 | -- | 3410.8 | 17.3 | 100 | Fig. S3b |
| DPPC+40% chol | 34.4 ± 27.1 | 0.80 ± 0.37 | -- | 4098.6 | 17.5 | 100 | Fig. S3b |

| Single-phase lipid sample | Polar angle, θ (deg) | Solid angle, Ω (π sr) | Phase-index (arb. units) | SM brightness (photon) | Background (photon/pixel) | Exposure time (ms) | Figure |
|---|---|---|---|---|---|---|---|
| DOPC | 28.0 ± 22.1 | 0.92 ± 0.33 | -- | 1770.6 | 5.0 | 30 | Fig. S3c |
| DOPC+40% chol | 32.9 ± 25.5 | 1.00 ± 0.32 | -- | 1720.8 | 5.2 | 30 | Fig. S3c |

Table S4. SMOLM orientation spectra statistics and experimental conditions for single-phase lipid samples imaged using the Duo-spot PSF.

| Single-phase lipid sample | Polar angle, θ (deg) | Solid angle, Ω (π sr) | Phase-index (arb. units) | SM brightness (photon) | Background (photon/pixel) | Exposure time (ms) | Figure |
|---|---|---|---|---|---|---|---|
| *Nile red (NR)* | | | | | | | |
| SPM | 46.9 ± 17.2 | 0.64 ± 0.41 | -- | 1617.9 | 12.7 | 100 | Fig. S10a |
| SPM+40% chol | 19.4 ± 14.1 | 0.18 ± 0.29 | -0.070 ± 0.302 | 2997.8 | 13.8 | 100 | Fig. S10a, S13b |
| DOPC | 41.5 ± 8.5 | 0.46 ± 0.40 | 0.069 ± 0.258 | 1265.2 | 2.4 | 20 | Fig. S13b |
| DOPC+40% chol | 42.8 ± 8.0 | 0.41 ± 0.39 | 0.080 ± 0.284 | 1264.7 | 2.3 | 20 | Fig. S13b |
| SPM+cer | 37.5 ± 15.7 | 0.71 ± 0.37 | 0.042 ± 0.271 | 1382.2 | 13.6 | 100 | Fig. S13b |

### Supplementary Movies

Movie S1. Raw SMLM images (43 successive frames, total 215 ms) of MC540 in a lipid mixture of DOPC/DPPC/chol (35:35:30, molar ratio). The gray lines show the boundary between Lo and Ld domains. The red box indicates the field of view used in Fig. 2b of the main text. The green dots indicate MC540 localizations with long binding times and confined lateral diffusion that contribute to the high localization counts within Lo domains (pixels with dramatically more localizations along the green line 1 in Fig. 2b(i)) and the local spikes in the SMLM line profiles (Fig. 2d(i), marked by asterisk). Red crosses represent MC540 localizations with high lateral diffusion within Ld domains.

Movie S2. SMOLM imaging of Nile red in SPM and SPM+40% chol SLBs using the Tri-spot PSF and Duo-spot PSF. For each PSF, fifty representative frames of raw images are shown in the left panel (red: x-polarized and blue: y-polarized fluorescence). Yellow crosses represent the estimated position of each molecule. Accumulated orientations (polar angle $\theta$) and wobble (solid angle $\Omega$) angles of single Nile red molecules are reported in the right panel. Scale bar: 2 μm.
